# FUS Oncofusion Protein Condensates Recruit mSWI/SNF Chromatin Remodelers via Heterotypic Interactions Between Prion-like Domains

**DOI:** 10.1101/2021.04.23.440992

**Authors:** Richoo B. Davis, Taranpreet Kaur, Mahdi Muhammad Moosa, Priya R. Banerjee

## Abstract

Fusion transcription factors generated by genomic translocations are common drivers of several types of cancers including sarcomas and leukemias. Oncofusions of the FET (FUS, EWSR1, and TAF15) family of proteins result from fusion of the prion-like domain (PLD) of FET proteins to the DNA-binding domain (DBD) of certain transcription regulators and are implicated in aberrant transcriptional programs through interactions with chromatin remodelers. Here, we show that FUS-DDIT3, a FET oncofusion protein, undergoes PLD-mediated phase separation into liquid-like condensates. Nuclear FUS-DDIT3 condensates can recruit essential components of the global transcriptional machinery such as the chromatin remodeler SWI/SNF. The recruitment of mammalian SWI/SNF is driven by heterotypic PLD-PLD interactions between FUS-DDIT3 and core subunits of SWI/SNF, such as the catalytic component BRG1. Further experiments with single-molecule correlative force-fluorescence microscopy support a model wherein the fusion protein forms condensates on DNA surface and enrich BRG1 to activate transcription by ectopic chromatin remodeling. Similar PLD-driven co-condensation of mSWI/SNF with transcription factors can be employed by other oncogenic fusion proteins with a generic *PLD*-*DBD domain* architecture for global transcriptional reprogramming.

## Introduction

FET fusion proteins are key drivers of several types of cancers including sarcomas and leukemias [1, 2]. These chimeric proteins are created by the oncogenic fusion of two non-homologous genes [3–5]. In the case of FET (FUS, EWSR1, and TAF15) family fusion proteins, their N-terminal disordered domain fuse to the DNA-binding domain of the transcription factor family ETS (E-twenty-six). The N-terminal domain of FET proteins features a low complexity sequence enriched in aromatic and polar amino acids (Q/N/Y/S/G) and is classified as ‘prion-like’ [6]. Prion-like domains (PLDs) are present in nearly 1% of the human proteome, predominantly in ribonucleoproteins (RNPs) including the FET proteins, TDP43, TIA1, and hnRNPA1 [7, 8]. PLD-containing proteins are highly enriched in various biomolecular condensates such as stress granules and transcription factor condensates [7, 9–11]. At the molecular level, PLDs enable the formation of dynamic protein condensates through a physical process known as liquid-liquid phase separation (LLPS), which is mediated by multivalent self-interactions among PLD chains involving distributed aromatic residues [12].

Although the phase separation of PLDs has been well characterized in the context of RNA binding proteins [13], the impact of PLDs fused to DNA-binding proteins such as transcription factors is relatively less explored. Transcription factors typically utilize their DNA-binding domain (DBD) to bind to specific gene loci and use their activation domains to recruit additional regulatory coactivators and RNA Polymerase II [14–17]. For FET oncofusion transcription factors, PLDs can act as the activation domain providing additional functionalities to the fusion transcription factors including the capacity to form phase-separated transcriptional hubs [18, 19]. Moreover, these fused PLDs may establish new interactions or modulate interactions with existing partners of the oncofusion transcription factor.

Recent studies have provided evidence that the PLDs of FET fusion oncoproteins interact with the mammalian SWI/SNF (mSWI/SNF) chromatin remodeling complex [2, 20–22]. The mSWI/SNF or BAF is an evolutionarily conserved multiprotein complex that uses ATP hydrolysis to reposition nucleosomes and remodel chromatin landscape [23–25]. The mSWI/SNF represents a wide variety of complexes with varying subunit compositions that are expressed in a developmental stage and tissue-specific manner [26, 27]. Predominantly localized at enhancers and promoters, mSWI/SNF plays key roles in regulating transcription [28]. Mutations in subunits of this complex are documented in ~ 20% of all cancers implying a high propensity for oncogenesis in the event of their dysregulation and/ or mislocalization [20, 23, 29, 30]. Since the subunits of mSWI/SNF lack DNA-recognition motifs, their recruitment to specific genomic locations is typically mediated via their interactions with transcription factors and coactivators [28, 31, 32].

In this study, we use the FET oncofusion FUS-DDIT3 as a model system to investigate the molecular behavior of such fusion oncoproteins and the mechanism of their engagement with the mSWI/SNF chromatin remodeler. FUS-DDIT3 (Type II fusion [33]; Fig. 1a) is composed of the PLD of FUS fused to the ETS family transcription factor, DDIT3 (see Table B: Materials & Methods for protein sequence), and is detected in more than 90% of the myxoid/round cell liposarcoma [34, 35]. Our results suggest that the PLD drives robust phase separation of FUS-DDIT3 in the mammalian cell nucleus, which is unlike the behavior of the parent protein FUS that remains predominantly soluble due to its interactions with nuclear RNAs. We further uncover the existence of PLDs in multiple subunits of the mSWI/SNF complex that can enable a synergistic engagement with the FET fusion oncoproteins through heterotypic PLD-PLD interactions. Such interactions provide a molecular mechanism for the recruitment of the mSWI/SNF to FET-oncoprotein condensates and reveal a molecular pathway for rewiring transcriptional programs by aberrant chromatin remodeling.

**Figure 1.**
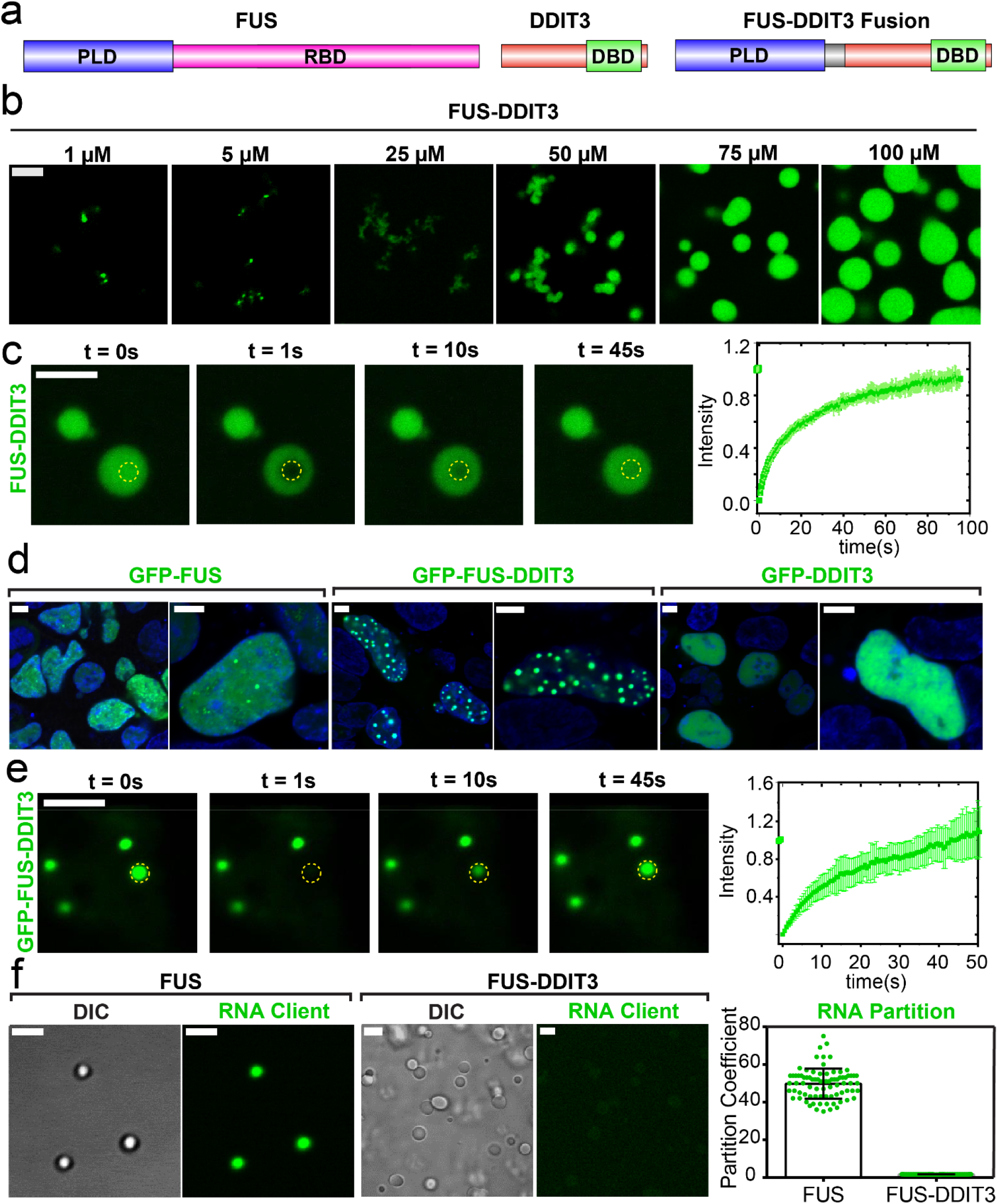
Oncofusion protein FUS-DDIT3 undergoes liquid-liquid phase separation *in vitro* and in mammalian cells. **(a)** Schematics of FUS, DDIT3, and FUS-DDIT3 domain architectures (PLD: Prion-like domain; RBD: RNA-binding domain; DBD: DNA-binding domain). **(b)** Fluorescence microscopy images of the assemblies formed by recombinantly purified FUS-DDIT3 (mixed with 250 nM AlexaFluor488-labeled FUS-DDIT3) at varying protein concentrations. **(c)** Representative fluorescence recovery after photobleaching (FRAP) images of recombinant FUS-DDIT3 condensates at 50 μM (t=0 s: pre-bleach; t=1 s: bleach; t>1 s: recovery). *Right:* The FRAP curve shows the average intensity and standard deviation of the intensity profiles over time (n=3). **(d)** Fluorescence microscopy images of HEK293T cells expressing GFP-tagged proteins (FUS, FUS-DDIT3, or DDIT3), as indicated. Hoechst was used to stain the nucleus and is shown in blue. **(e)** Representative fluorescence recovery after photobleaching (FRAP) images of GFP-FUS-DDIT3 condensates expressed in HEK293T cells (t=0 s: pre-bleach; t=1 s: bleach; t>1 s: recovery). *Right:* The FRAP curve shows the average intensity and standard deviation of the intensity profiles over time (n=3). **(f)** Partitioning of FAM-labeled RNA client into condensates of recombinant FUS-DDIT3 (50 μM) and recombinant FUS (6 μM). (DIC: Differential Interference Contrast; FAM: 6-Carboxyfluorescein). *Right:* Enrichment is calculated as partition coefficients. Mean and standard deviation are shown. The scale bar is 5 μm for all images.

## Results

### FUS-DDIT3 undergoes LLPS in vitro and in mammalian cells

Prior studies have established a key role of the prion-like domain (PLD) in driving the phase separation of FUS into liquid-like condensates [13, 36, 37]. We, therefore, wanted to test if the PLD enables FUS-DDIT3 fusion protein to undergo a similar liquid phase condensation. Utilizing recombinant FUS-DDIT3, we found that the fusion oncoprotein forms condensates *in vitro* in a concentration-dependent manner (Fig. 1b). FUS-DDIT3 condensates were stable across a broad range of NaCl concentrations (10-300 mM, Fig. S1a), and the presence of polymer crowders facilitated their formation (Fig. S1b). Fluorescence recovery after photobleaching (FRAP) experiments showed that FUS-DDIT3 molecules within these condensates are dynamic, implying a liquid-like behavior (Fig. 1c). Interestingly, we observed that FUS-DDIT3 assemblies appear as a mixture of “irregular” bodies and spherical condensates at lower protein and crowder concentrations but form predominantly large spherical condensates at higher concentrations (C_FUS-DDIT3_ > 50 µM and Ficoll PM70 ≥ 10%; Figs.1b & S1b). This is analogous to the previously reported study on SPOP-DAXX condensates where competition between inter-chain interactions of variable strengths determine the material state of the resultant assemblies [38, 39]. When compared with the PLD of FUS alone, we observed that the FUS-DDIT3 fusion protein formed condensates at a significantly lower protein concentration (Fig.1b & S1c). These data suggest that DDIT3 may directly contribute to the homotypic interactions between FUS-DDIT3 molecules. Consistent with this idea, we observed that recombinant DDIT3 itself can form micron-scale assemblies *in vitro* that appear gel-like and are less dynamic than FUS-DDIT3 assemblies (Figs. S1d&e). Thus, intermolecular PLD-PLD interactions along with DDIT3-DDIT3 interactions synergistically facilitate the formation of FUS-DDIT3 condensates.

To investigate the phase behavior of FUS-DDIT3 in cell culture models, we expressed GFP-tagged FUS-DDIT3 in HEK293T cells and observed the formation of FUS-DDIT3 enriched spherical assemblies in the nucleus (Fig. 1d, center panel). These FUS-DDIT3 foci were dynamic as evidenced by their near-complete fluorescence recovery after photobleaching (Fig. 1e). We also observed that FUS-DDIT3 droplet/foci formation was concentration-dependent as cells with low concentrations of transgenically expressed FUS-DDIT3 lacked these foci (Fig. S1f), suggesting that the oncofusion protein’s condensation is mediated via an LLPS mechanism. On the contrary, both GFP-DDIT3 and GFP-FUS showed a diffused distribution in the nucleus, with small punctate structures observed for GFP-FUS (Fig. 1d). Taken together, these results demonstrate that the PLD of FUS when fused to the transcription factor DDIT3 can drive phase separation of the fusion protein both *in vitro* and in the cell nucleus.

### FUS-DDIT3 forms stable condensates within the RNA-rich nuclear environment

To understand the distinct behavior of FUS-DDIT3 and the parent FUS protein in the nucleus, we turned to the role of RNA in regulating protein phase separation. FUS is an RNA binding protein, and RNA can regulate the phase behavior and material properties of FUS condensates [40, 41]. Although recombinant FUS can undergo phase separation at low protein concentrations (~2 μM) *in vitro*, which is lower than FUS’s nuclear concentration, the high level of RNA in the nucleus suppresses phase separation of FUS [42]. Our previous studies have established that RNA has a dual role in regulating the LLPS of FUS protein. At low RNA concentrations, RNA promotes FUS phase separation via the formation of “sticky” complexes, whereas at higher concentrations, it inhibits FUS condensation due to the formation of negatively charged FUS-RNA complexes [41]. Since FUS-DDIT3 lacks an RNA-binding domain, we hypothesized that FUS-DDIT3 may have a lower affinity for RNA and therefore, FUS-DDIT3 condensates may be refractory to the high RNA concentrations within the nucleus. This idea is directly supported by our observations that nuclear FUS-DDIT3 forms stable condensates whereas nuclear FUS remains predominantly diffused (Fig. 1d). To test this model further, we examined the partitioning of a short RNA client ([6FAM]UGAAGGAC) into recombinant FUS and FUS-DDIT3 condensates *in vitro*. While FUS condensates could readily interact with and enrich RNA (partition coefficient: 50±8; Fig. 1f), FUS-DDIT3 condensates did not show any significant enrichment (partition coefficient: 1.60±0.03; Fig. 1f). We also note a subset of GFP-FUS-DDIT3 condensates exhibited hollow spherical morphologies in the nucleus (Fig. S2) similar to what was observed previously for the condensates formed by TDP43 mutants with impaired RNA binding ability [43]. Overall, these results indicate that RNA does not significantly interact with FUS-DDIT3 and provide an explanation for FUS-DDIT3’s ability to form robust condensates within the RNA-rich environment of the cell nucleus where FUS primarily remains soluble.

### FUS-DDIT3 condensates enrich BRG1, a catalytic subunit of the mSWI/SNF complex

Nuclear condensates can compartmentalize machineries responsible for many biochemical events such as transcription, splicing, and chromatin organization [44]. Recent studies have indicated that transcription factor condensates can activate genes by creating phase-separated transcriptional hubs at super-enhancer sites that enrich coactivators and RNA polymerase II [45–47]. Based on our observations that FUS-DDIT3 forms stable nuclear condensates, we aimed to investigate if these condensates can enrich transcriptional activators/coactivators. Previous studies have established that FET fusion protein interactomes are enriched in the components of mSWI/SNF complex and that BRG1, the key ATPase subunit that drives chromatin remodeling by mSWI/SNF, co-immunoprecipitates with the FUS-DDIT3 fusion protein [2, 22]. Therefore, we hypothesized that nuclear FUS-DDIT3 condensates may compartmentalize BRG1. To test this idea, we expressed GFP-tagged BRG1 along with mCherry-tagged FUS-DDIT3 in HEK293T cells. We observed that BRG1 remained diffused in the nucleus in absence of FUS-DDIT3, but was readily recruited within FUS-DDIT3 nuclear condensates when co-expressed (Fig. 2a-b&S3). Independently, using purified proteins *in vitro*, we observed that recombinant BRG1 protein is enriched within the FUS-DDIT3 condensates [partition coefficient: 27±7 (Fig. 2c)]. These data suggest that phase separation of the FUS-DDIT3 leads to the enrichment and ectopic compartmentalization of the chromatin remodeler BRG1 into nuclear oncoprotein condensates.

**Figure 2.**
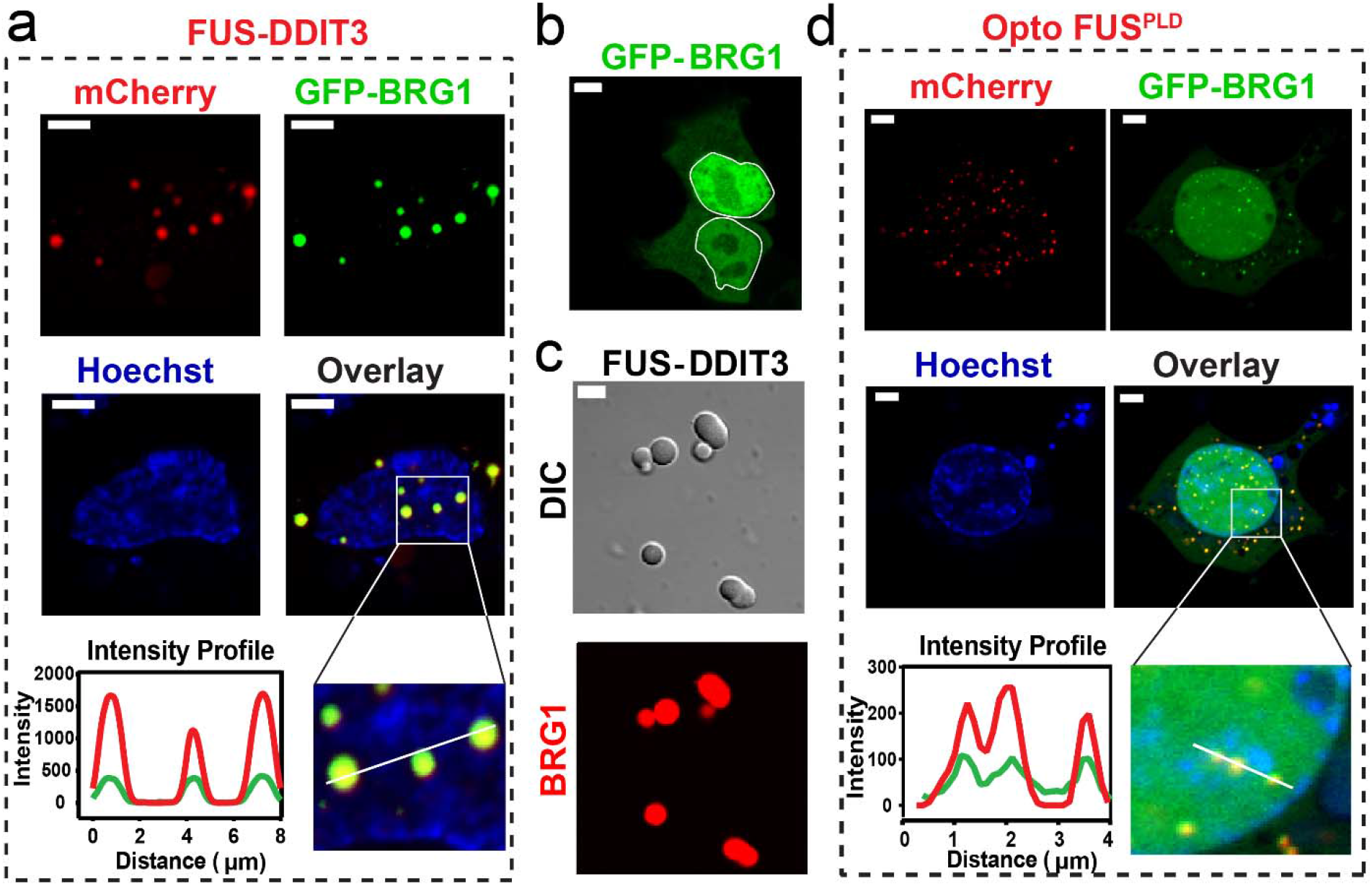
FUS-DDIT3 condensates enrich mSWI/SNF subunit BRG1 via the FUS^prion-like domain^. **(a)** HEK293T cells co-expressing GFP-BRG1 and Cry2-mCherry-FUS-DDIT3. FUS-DDIT3 condensates were formed via protein overexpression without any blue-light activation and co-localized with GFP-BRG1 (see Materials & Methods). Hoechst (blue) was used to stain the nucleus. The region demarcated in the white square is magnified and the fluorescence intensity profiles are shown across the linear section (white line). Green represents the intensity profile of GFP-BRG1 and red represents the profile for Cry2-mCherry-FUS-DDIT3. **(b)** HEK293T cells expressing the GFP-BRG1 protein show diffused distribution in both nucleus (demarcated by the white line) and the cytoplasm. **(c)** Partitioning of RED-tris-NTA-labeled BRG1 (0.5 μM) into recombinant FUS-DDIT3 condensates (50 μM). **(d)** HEK293T cells co-expressing GFP-BRG1 and Cry2-mCherry-FUS^PLD^ (OptoFUS^PLD^). OptoFUS^PLD^ droplets were formed by activating with blue light for sixty seconds and then enrichment of GFP-BRG1 was analyzed within these light-induced condensates. Hoechst (blue) was used to stain the nucleus. The region demarcated in the white square is magnified and the fluorescence intensity profiles are shown across the linear section (white line). Green represents the intensity profile of GFP-BRG1 and red represents the profile for Cry2-mCherry-FUS^PLD^. The scale bar is 5 μm for all images.

What is the molecular mechanism of BRG1 recruitment within FUS-DDIT3 condensates? Linden *et al.* reported that the N-terminal prion-like domains of FET proteins can coimmunoprecipitate with multiple subunits of the mSWI/SNF complex [2]. In addition to the PLD-containing FET oncofusions, PLD-harboring transcription activators such as EBF1 and MN1 have also been suggested to interact with BRG1 [10, 48]. We, therefore, hypothesized that the PLD of FUS-DDIT3 is responsible for recruiting BRG1 within FUS-DDIT3 condensates. To test this, we used a previously characterized OptoFUS^PLD^ construct containing a Cry2 tag that homo-oligomerizes on exposure to blue light (488 nm) and nucleates formation of the FUS^PLD^ condensates. [49]. We observed that when co-expressed with GFP-BRG1 protein, OptoFUS^PLD^ droplets can enrich GFP-BRG1 (Fig. 2d & S3). These results suggest that the PLD of FUS is sufficient to recruit and compartmentalize the chromatin remodeler BRG1 in FUS-DDIT3 condensates.

### Prion-like domains can act as scaffolds to recruit mSWI/SNF proteins in FUS-DDIT3 condensates

PLDs are known to self-assemble into a variety of assemblage including phase-separated condensates, amorphous aggregates, and fibrillar solids [13, 36, 37]. Furthermore, distinct PLDs are also known to cooperate and co-condense through heterotypic PLD-PLD interactions in transcription factor condensates [11]. Given the PLD of FUS is sufficient to recruit BRG1 into FUS-DDIT3 condensates, we enquired if BRG1 also carries a prion-like domain. Sequence analysis using the PLAAC algorithm [50] revealed that the N-terminus of BRG1 (aa 1-340) is disordered and has a significant prion-like amino acid composition (Fig. S4). Therefore, it is conceivable that heterotypic PLD^FUS^-PLD^BRG1^ interactions can recruit BRG1 into FUS-DDIT3 condensates. We first characterized the phase separation capability of BRG1^PLD^ and observed that recombinant BRG1^PLD^ has a low intrinsic tendency to phase separate *in vitro*, only forming condensates at relatively high protein and crowder concentrations (Fig. S5a). Furthermore, when expressed in HEK293T cells, OptoBRG1^PLD^ proteins did not form condensates even at concentrations that are ~ 4-fold higher than that required for the formation of OptoFUS^PLD^ condensates (Figs. 3a and S5b). Thus, the N-terminal prion-like domain of BRG1 has a weak capacity to undergo self-condensation both *in vitro* and in cells.

**Figure 3.**
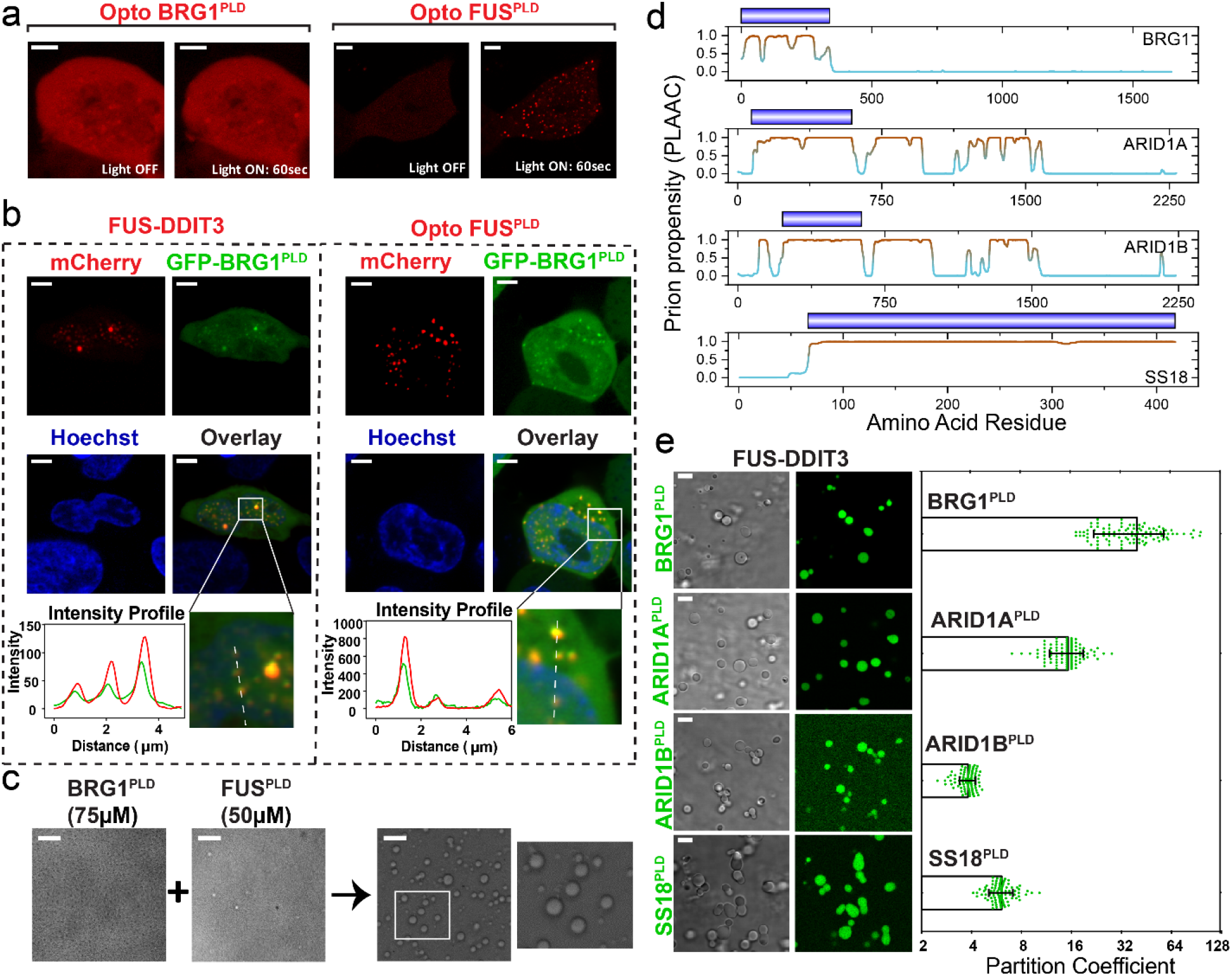
Many mSWI/SNF complex subunits contain PLDs and enrich into FUS-DDIT3 condensates via heterotypic PLD-PLD interactions. **(a)** HEK293T cell expressing OptoBRG1^PLD^ (*left*) or OptoFUS^PLD^ (*right*) before and after activation with blue light for 60 seconds. Scale bar is 5 μm **(b)** HEK293T cells co-expressing GFP-BRG1^PLD^ and either Cry2-mCherry-FUS-DDIT3 (*left*) or OptoFUS^PLD^ (*right*). OptoFUS^PLD^ droplets were formed by blue light stimulation for sixty seconds and then enrichment of GFP-BRG1^PLD^ was analyzed within the condensates. Cry2-mCherry-FUS-DDIT3 condensates were formed via protein overexpression without any blue-light activation. Hoechst (blue) was used to stain the nucleus. The region demarcated in the white square is magnified and the fluorescence intensity profile is shown across the linear section (white line). Green represents the intensity profile of GFP-BRG1^PLD^ and red represents the profile for either Cry2-mCherry-FUS-DDIT3 or OptoFUS^PLD^. The scale bar is 5 μm. **(c)** Co-condensation of purified BRG1^PLD^ and FUS^PLD^. No droplets were observed before mixing. The region demarcated in the white square is magnified and shown for better clarity. The scale bar is 10 μm. **(d)** PLAAC analysis showing multiple regions with high prion-propensity for the four selected subunits of mSWI/SNF complex (see Fig. S4 for PLAAC profiles for all PLD-containing mSWI/SNF complex subunits). Domains corresponding to royal blue bars were recombinantly expressed and purified and used in our experiments (*panel e*). **(e)** Partitioning of recombinant PLDs from *panel d* (fluorescently labeled with AlexaFluor488) into FUS-DDIT3 condensates (50 μM). Enrichment is calculated as partition coefficient. Mean and standard deviation are shown. (Partition coefficients: BRG1^PLD^ = 40±20; ARID1A^PLD^ = 15±4; ARID1B^PLD^ = 3.8±0.4; SS18^PLD^ = 6±1). The scale bar is 10 μm.

Although the PLD of BRG1 did not undergo phase separation in cells, we tested whether it could engage with the PLD of FUS through heterotypic PLD-PLD interactions. We expressed GFP-tagged BRG1^PLD^ in cells expressing OptoFUS^PLD^. Upon blue light activation, cells expressing OptoFUS^PLD^ formed condensates of FUS^PLD^ that enriched GFP-BRG1^PLD^ (Fig. 3b right panel). Using independent experiments, we further observed that GFP-BRG1^PLD^ is enriched within the nuclear FUS-DDIT3 condensates (Fig. 3b left panel). Consistent with these results, we also found that when recombinant BRG1^PLD^ and FUS^PLD^ were mixed at a concentration below their individual phase separation thresholds *in vitro*, they form co-condensates (Fig. 3c and S6). This is likely due to heterotypic PLD-PLD interactions leading to the lowering of phase separation saturation concentration of the PLD mixture [51]. Thus, BRG1^PLD^ can interact and undergo co-condensation with the prion-like domain of FUS.

A broader analysis using PLAAC revealed that in addition to BRG1, multiple subunits of the mSWI/SNF complex contain *bona fide* PLDs (Fig. 3d and S4). These include SMARCC1 and SMARCC2, which along with BRG1, are three of the four core subunits of the complex that orchestrate chromatin remodeling [52, 53]. In addition, accessory (and/or signature) subunits such as ARID1A and ARID1B, mutations of which are highly correlated with oncogenic transformation [23, 54] also have long tracts of prion-like sequences (Fig. 3d and S4). To test if these PLDs could also mediate interactions with FET fusion oncoproteins, we reconstituted FUS-DDIT3 condensates *in vitro* and looked for enrichment of fluorescently labeled PLDs from the three associated factors with the longest PLDs: ARID1A, ARID1B, and SS18. All of the tested PLDs were enriched within FUS-DDIT3 condensates (Fig. 3e), although with varied partition coefficients (partition coefficient ranges from ~ 4 to 40), suggesting different strengths of their interactions. This can be attributed to distinct sequence compositions, chain lengths, and charge patterns of the tested PLDs. Together, these data suggest that the prion-like domain of FET fusion oncoproteins can engage with prion-like domains of multiple mSWI/SNF proteins and recruit them to ectopic nuclear condensates.

### FUS-DDIT3 forms condensates on dsDNA and recruits chromatin remodeler BRG1

Our results discussed above suggest that the PLD of FUS enables phase separation of the DDIT3 transcription factor. Transcription factors can interact with the surface of DNA and such interactions can contribute to their phase separation at specific genomic loci [19, 55, 56]. To test if FUS-DDIT3 can form condensates on DNA, we tethered a single double-stranded (ds) λ-phage genomic DNA between two optically trapped polystyrene beads using laminar-flow in a microfluidic glass chamber (see Materials & Methods for further details and Figs. 4a&S7). Upon transferring this tethered DNA molecule into a channel containing 250 nM AlexaFluor488-labeled FUS-DDIT3, we detected the formation of distinct FUS-DDIT3 clusters on the dsDNA (Fig 4b, top left panel). The FUS-DDIT3 clusters formed at multiple loci on a single DNA chain. This is likely because FUS-DDIT3 interacts promiscuously with the λ-phage genomic DNA [19]. Next, to determine if these FUS-DDIT3 condensates can recruit BRG1, the single-molecule DNA tether with FUS-DDIT3 condensates was transferred to a microfluidic channel containing 10 nM of RED-tris-NTA-labeled BRG1 (Fig. S7). In line with our above-mentioned results, we observed that the BRG1 co-localizes with the FUS-DDIT3 condensates on the surface of dsDNA (Fig. 4b, top right panel). The co-localization of BRG1 within FUS-DDIT3 condensates was confirmed using the confocal images as well as by analyzing the intensity profiles which revealed a clear overlap of FUS-DDIT3 and BRG1 peaks (Fig. 4b bottom panel). We also noted a few isolated BRG1 peaks that are present across the dsDNA without any detectable spatial overlap with FUS-DDIT3 condensates and the occurrence of such peaks increases with increasing bulk concentration of BRG1 from 10 nM to 50 nM (Fig. S8). Such foci formation may indicate the presence of a possible interaction between BRG1 and the λ-phage genomic DNA that is independent of FUS-DDIT3.

**Figure 4.**
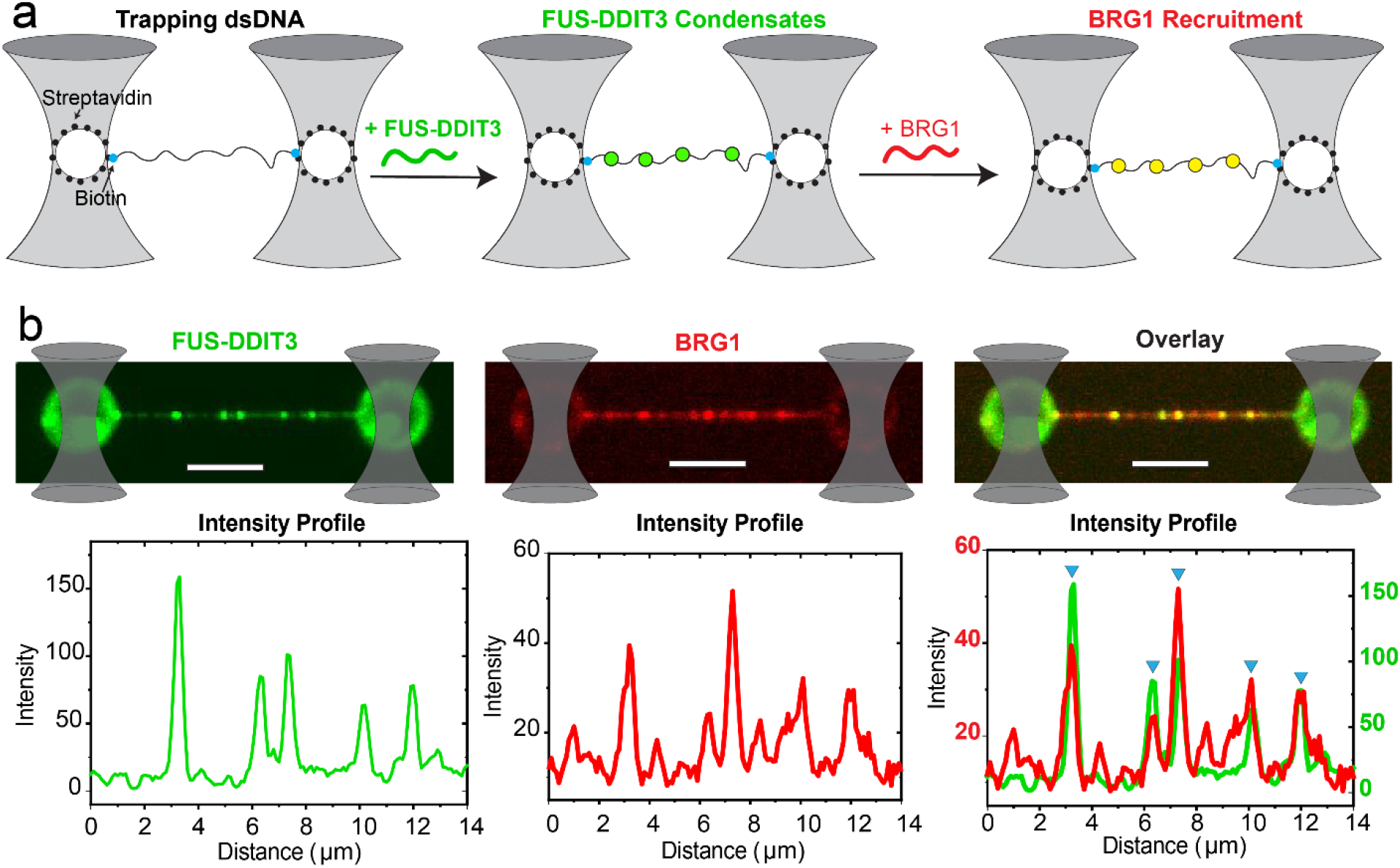
FUS-DDIT3 condensates on dsDNA recruit BRG1. **(a)** A schematic representation of the single-molecule DNA tethering assay. A single molecule of dsDNA (with ends biotinylated; represented by solid cyan circles) is tethered between two optically trapped polystyrene beads (traps are shown as grey cones; beads are shown as white circles) coated with streptavidin (solid black circles). The transfer of DNA molecule into a microfluidic channel containing FUS-DDIT3 leads to the formation of FUS-DDIT3 condensates (solid green circles). Subsequent transfer of the DNA chain decorated with FUS-DDIT3 condensates to a separate microfluidic channel containing BRG1 leads to the recruitment of BRG1 into the FUS-DDIT3 condensates (solid yellow circles). **(b)** Experimental data: multicolor confocal fluorescence micrographs and intensity profiles showing the formation of FUS-DDIT3 puncta/condensates (green) on a single DNA molecule followed by the recruitment of BRG1 (red) into the FUS-DDIT3 condensates. Blue triangles show the position of the overlapping intensity peaks of FUS-DDIT3 and BRG1 representing the recruitment of BRG1 into FUS-DDIT3 puncta/condensates. [FUS-DDIT3] = 250 nM and [BRG1] = 10 nM. The scale bar is 5 μm for all images.

### Many recurrent oncogenic translocations result in the fusion of prion-like domains with transcriptional regulators

Our results, presented so far, indicate that both partners in the fusion pair (*i.e.,* FUS^PLD^ and DDIT3) are likely to contribute to the neomorphic activity of the FET oncofusion protein – the DDIT3^DBD^ can recruit the fusion transcription factor to specific genomic loci, whereas the FUS^PLD^ can drive their condensation and the subsequent recruitment of the chromatin remodeler mSWI/SNF. Since many transcription factors are involved in oncogenic fusions [57], we asked whether fusion between the *DNA localization domain of transcriptional regulators* and *PLD of prion-containing proteins* represents a generic category of oncogenic transcription regulators. To answer this, we focused on translocations involving transcription regulators. Specifically, we retrieved sequences of transcription regulator fusions that are recurrently present in cancer patients using the following selection criteria: (*i*) present in at least 25 curated patient samples in the COSMIC database [58], and (*ii*) present in >10% of tumor samples in a given malignancy (The list of recurrent translocations involving transcriptional regulators was obtained from the reference [59]). We further narrowed down our list to the oncofusions that harbor at least one prion-like domain and identified a total of 16 different recurrent oncofusions involving 10 fusion pairs (Table 1). These proteins represent *in-frame* fusions between a prion-like domain and a specific DNA localization domain. The DNA localization domains were either *a specific DNA-binding domain* (*e.g.,* ERG^DBD^ in FUS-ERG fusion) or *a protein-protein interaction motif that specifically interacts with transcription regulators* (*e.g.,* SSX^SSXRD^ in SS18-SSX family fusions). In all these oncofusions, prion-like domains and DNA-localization domains originate from the two separate genes involved in respective *in-frame* genomic translocations. Not surprisingly, we find that many of these recurrent oncofusions have already been implicated in chromatin reorganization (Table 1). Therefore, as observed for FUS-DDIT3 fusion protein in our experiments, other PLD-containing oncogenic transcription regulators may similarly engage with mSWI/SNF chromatin remodelers to drive aberrant transcriptional outcomes via protein co-condensation.

**Table 1:**
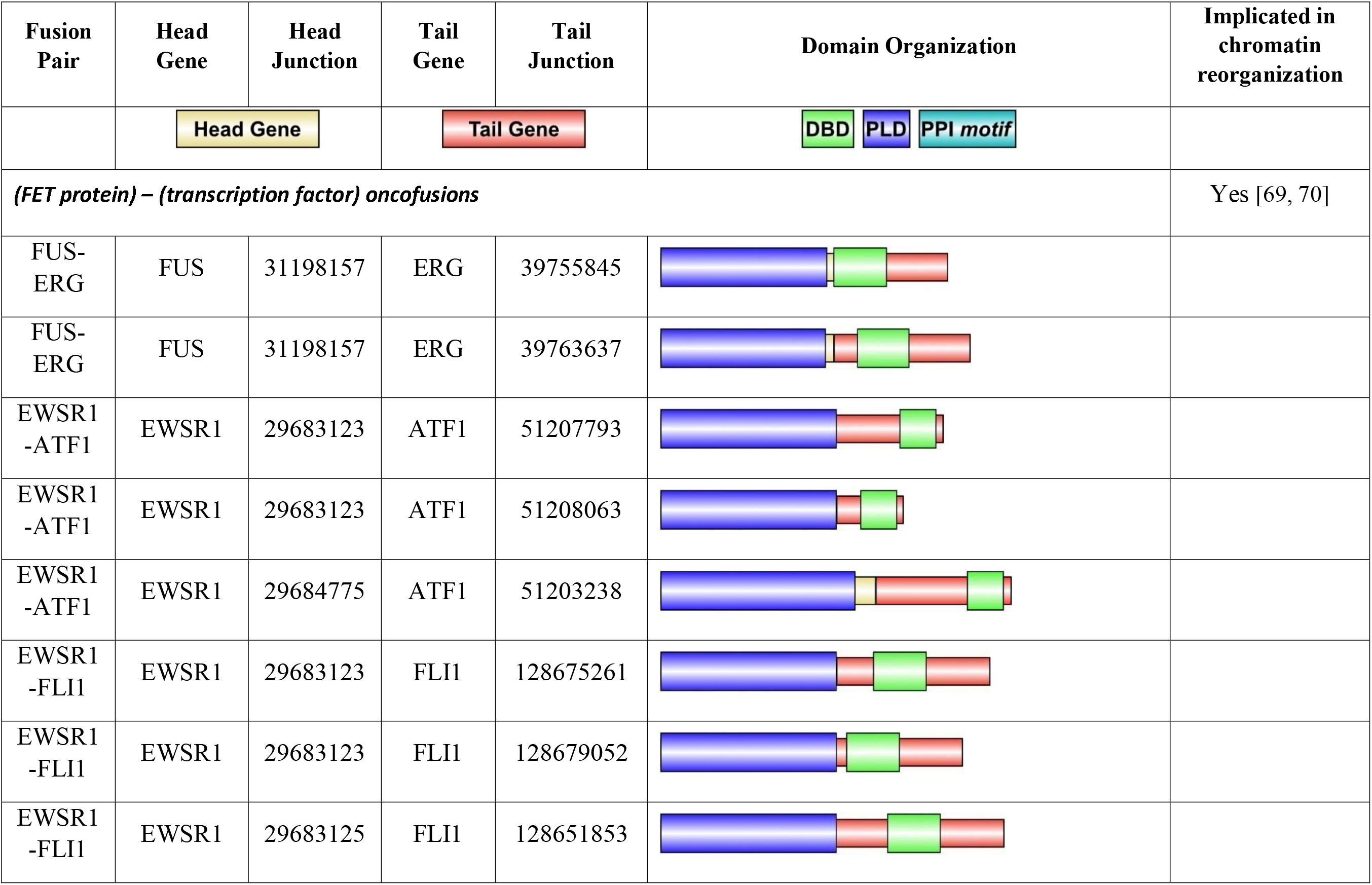
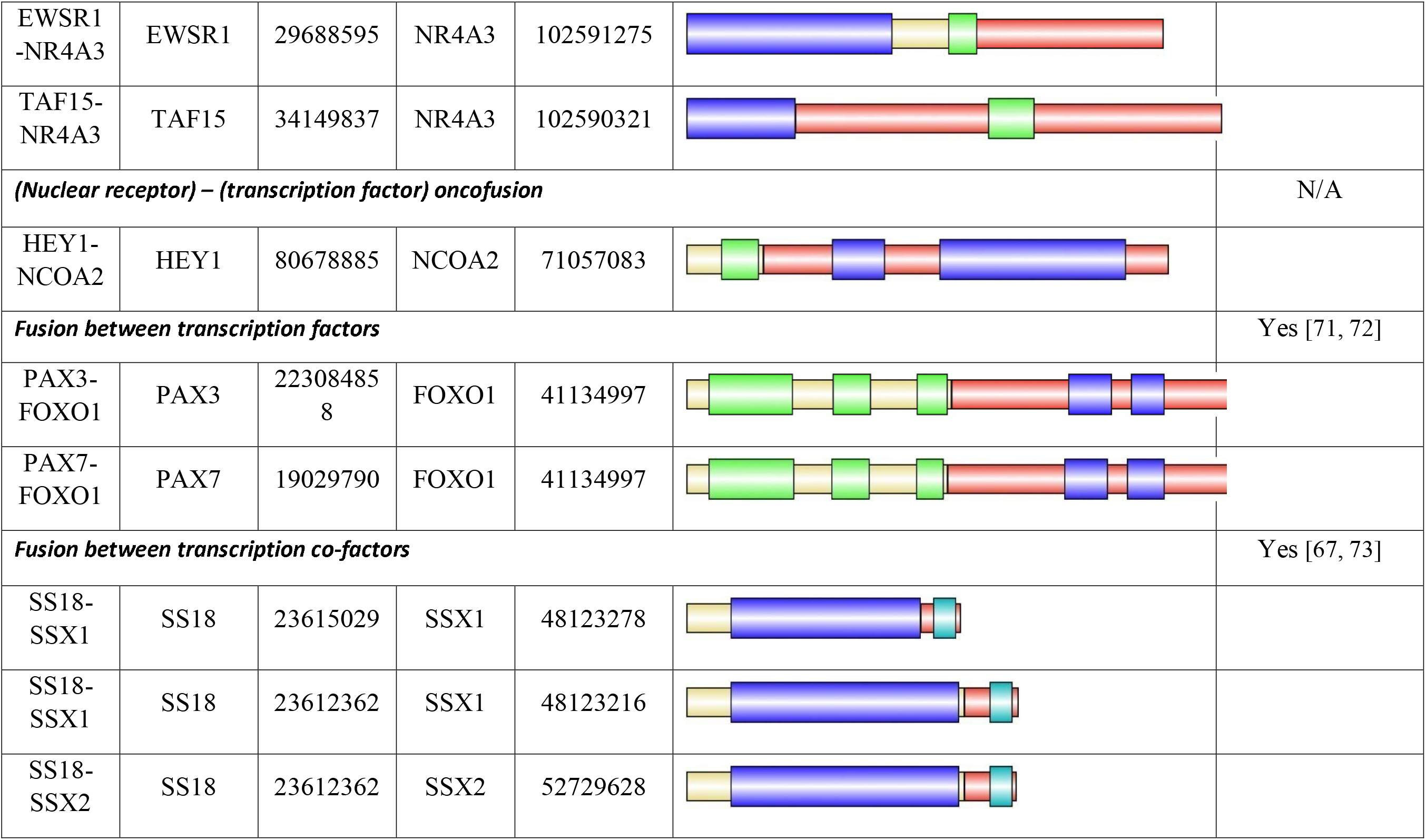
Recurrent fusions of transcriptional regulators with prion-like domains. Legends: DBD – DNA-binding domain; PLD – Prion-like domain; PPI motif – Protein-protein interaction motif.

| Fusion Pair | Head Gene | Head Junction | Tail Gene | Tail Junction | Domain Organization | Implicated in chromatin reorganization |
| --- | --- | --- | --- | --- | --- | --- |
|  | Head Gene |  | Tail Gene |  | DBD PLD PPI motif |  |
| *(FET protein) – (transcription factor) oncofusions* |  |  |  |  |  | Yes [69, 70] |
| FUS-ERG | FUS | 31198157 | ERG | 39755845 |  |  |
| FUS-ERG | FUS | 31198157 | ERG | 39763637 |  |  |
| EWSR1-ATF1 | EWSR1 | 29683123 | ATF1 | 51207793 |  |  |
| EWSR1-ATF1 | EWSR1 | 29683123 | ATF1 | 51208063 |  |  |
| EWSR1-ATF1 | EWSR1 | 29684775 | ATF1 | 51203238 |  |  |
| EWSR1-FLI1 | EWSR1 | 29683123 | FLI1 | 128675261 |  |  |
| EWSR1-FLI1 | EWSR1 | 29683123 | FLI1 | 128679052 |  |  |
| EWSR1-FLI1 | EWSR1 | 29683125 | FLI1 | 128651853 |  |  |
| EWSR1-NR4A3 | EWSR1 | 29688595 | NR4A3 | 102591275 |  |  |
| TAF15-NR4A3 | TAF15 | 34149837 | NR4A3 | 102590321 |  |  |
| ***(Nuclear receptor) – (transcription factor) oncofusion*** |  |  |  |  |  | N/A |
| HEY1-NCOA2 | HEY1 | 80678885 | NCOA2 | 71057083 |  |  |
| ***Fusion between transcription factors*** |  |  |  |  |  | Yes [71, 72] |
| PAX3-FOXO1 | PAX3 | 223084858 | FOXO1 | 41134997 |  |  |
| PAX7-FOXO1 | PAX7 | 19029790 | FOXO1 | 41134997 |  |  |
| ***Fusion between transcription co-factors*** |  |  |  |  |  | Yes [67, 73] |
| SS18-SSX1 | SS18 | 23615029 | SSX1 | 48123278 |  |  |
| SS18-SSX1 | SS18 | 23612362 | SSX1 | 48123216 |  |  |
| SS18-SSX2 | SS18 | 23612362 | SSX2 | 52729628 |  |  |

## Discussion

A major consequence of FET oncofusions is the transfer of the prion-like domain from an RNA-binding protein to a DNA-binding protein. Our results described here along with two recent reports suggest that FET-fusion oncoproteins can form ectopic nuclear condensates [18, 19]. These reports also suggest that FET-fusion protein condensates can activate transcription at DNA enhancer sites by engaging with essential transcriptional co-activators such as BRD4 and by recruiting RNA polymerase II [18, 19]. In our study, we focus on characterizing the interactions between FET-fusion condensates with the ATP-dependent chromatin remodeler complex mSWI/SNF. Chromatin remodeling is one of the primary and essential steps in the regulation of gene expression and the aberrant targeting of mSWI/SNF complexes can be detrimental to physiologic cellular processes [60]. Our results reveal that FUS-DDIT3 condensates can compartmentalize BRG1, a key catalytic subunit (ATPase) of mSWI/SNF that is responsible for ATP-dependent remodeling of DNA-histone interactions. Since a continuous activity of the mSWI/SNF is often required to maintain the appropriate chromatin state at the target genomic locus [61, 62], the selective enrichment of mSWI/SNF components by FUS-DDIT3 condensates provides two possible routes to transcriptional reprograming (Fig. 5). *First*, condensates of FET-fusion proteins formed at ectopic genomic regions, as specified by binding sites of the DNA-binding domain, can recruit BRG1 and modify local chromatin folding to activate transcription (Fig. 5, *left panel*). Previous studies with FET oncoproteins, such as EWS-FLI1 and FUS-DDIT3 showed retargeting of chromatin remodelers to microsatellites and enhancers respectively [20, 63], lending support to this model. Congruently, condensates formed by both FET proteins and FET fusion-oncoproteins are capable of enhancing gene transcription [19, 64]. *Alternatively*, we propose a model where sequestration of mSWI/SNF subunits away from their natural targets and trapping them in ectopic condensates can contribute to a loss-of-function phenotype (Fig. 5, *right panel*). This model is consistent with the recently reported observations that mSWI/SNF complex suppresses the H3K27Me3 histone modification under normal conditions, but when FET oncoproteins are expressed, the H3K27Me3 levels within the cells are upregulated likely due to the sequestration of mSWI/SNF away from its physiological target sites [2].

**Figure 5.**
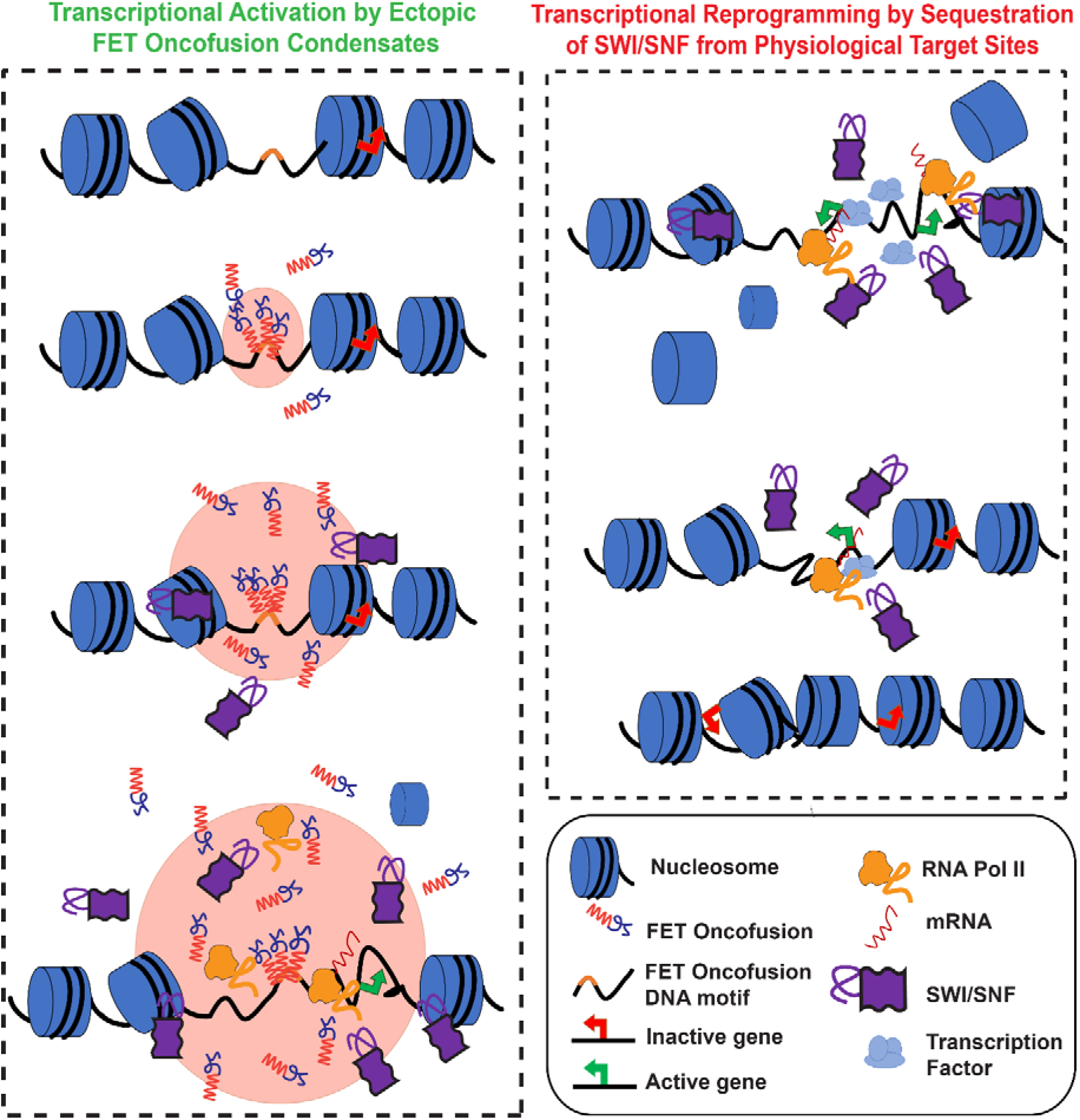
Proposed model for transcriptional reprogramming by FET-fusion oncoproteins. *Left:* FET fusion proteins can bind to specific DNA motifs defined by the DNA-binding domain at physiologically inactive genes to form phase-separated condensates. The process of LLPS is mediated by the prion-like domain. FET fusion condensates can recruit the ATP-dependent chromatin remodeler SWI/SNF leading to remodeling of the closed chromatin to an open chromatin state, which subsequently provides access to transcriptional coactivators and RNA polymerase II to mediate transcription of otherwise silenced genes. *Right:* The sequestration of chromatin remodeler SWI/SNF into FUS-DDIT3 condensates, as shown in the left panel, can lead to diminished chromatin remodeling activity at physiologically active genes where continuous SWI/SNF activity is required. This sequestration of SWI/SNF away from the physiological target genes may trigger a switch to a closed chromatin state, thereby decreasing their transcriptional output.

At the molecular level, the presence of PLDs in multiple subunits of the mSWI/SNF (Fig. S4) suggests that PLD-mediated interactions could be a generic mechanism for the recruitment of these complexes to their target sites. This is based on our observations that the PLD of FUS is sufficient to target the mSWI/SNF ATPase, BRG1, to FUS-DDIT3 condensates (Figs. 2&3). Moreover, the PLDs of mSWI/SNF proteins can synergistically engage with FET^PLD^ and reduce the phase separation threshold as observed in mixtures of FUS^PLD^ and BRG1^PLD^ (Fig. S6). These data also indicate that heterotypic PLD-mediated co-condensation may play a fundamental role in the functional assembly of the mSWI/SNF complex itself.

We envision that hijacking of mSWI/SNF complex by FET oncofusions through heterotypic PLD-PLD interactions is a generic strategy employed by many aberrant transcription regulators. This is supported by our analysis that many recurrent oncogenic translocations involve the fusion of a prion-like domain to a DNA recruitment domain and result in a widespread reorganization of the chromatin landscape (Table 1). In addition to the FET oncofusions, SS18-SSX fusions represent an interesting category of oncogenic translocations. In SS18-SSX fusions, the prion-like domain of SS18 is fused to the C-terminal segments of SSX family proteins containing the SSXRD domain [65, 66]. SSXRD domain is a protein-protein interaction module that directly interacts with the polycomb complex DNA-binding protein, KDM2B [67]. Not surprisingly, SS18-SSX fusions recruit mSWI/SNF complex to KDM2B-binding genomic loci and result in the aberrant activation of *numerous* otherwise repressed genes [67, 68]. Therefore, similar to the FUS^PLD^, it is likely that SS18^PLD^ is capable of recruiting mSWI/SNF chromatin remodelers via SS18^PLD^–SWI/SNF^PLD^ interactions. However, unlike FET fusions where the neomorphic transcriptional activator is recruited to specific genomic locations via DNA-binding domains fused to the FET^PLD^, SS18-SSX fusions may utilize specific protein-protein interaction module from SSX family members (*i.e,* the SSXRD domain) to recruit oncofusion proteins to specific loci.

In summary, our results provide a molecular mechanism with regards to how FET oncofusions can synergistically engage with the ATP-dependent chromatin remodeler mSWI/SNF. Our bioinformatics analysis reveals that PLD fusions to DNA-binding domains go beyond FET fusion oncoproteins and the proposed role of heterotypic PLD-PLD interactions in recruiting mSWI/SNF complex at non-native genomic loci may play a central role in all such cases. Future studies can test the generality of this model and subsequently target the phase separation and/or mSWI/SNF complex engagement capacities of PLD containing oncofusion proteins for potential cancer therapeutics.

## Data Availability

All data relevant to the findings of this manuscript are included in the manuscript and the supplementary appendix. Additional data are available from the authors upon reasonable request.

## Acknowledgment

The authors acknowledge Dr. Sarah Walker of the University at Buffalo, SUNY for valuable help with protein purification. P.R.B. acknowledges the College of Arts and Sciences at the University at Buffalo, SUNY, and the National Institute of General Medical Sciences (NIGMS) of the National Institutes of Health (R35 GM138186) for financial support.

## Author Contributions

P.R.B. and M.M.M. conceived the idea. P.R.B., M.M.M., R.D., and T.K. designed the study. R.D. and T.K. performed the experiments and data analysis. M.M.M. performed the bioinformatics analysis. All authors contributed to writing the manuscript.

