## Supplementary Information for "FUS Oncofusion Protein Condensates Recruit mSWI/SNF Chromatin Remodelers via Heterotypic Interactions Between Prion-like Domains"

### Materials & Methods

**Protein expression, purification, and labeling:** A list of proteins used in the study is provided in Table A and their amino acid sequence is provided in Table B. Codon optimized proteins of interest were gene-synthesized by GenScript USA Inc. (Piscataway, NJ, USA). The plasmid vector was a gift from Scott Gradia (Addgene plasmid # 29706). Proteins were expressed, purified, and fluorescently labeled as described in our earlier work with one modification [1]. Cells were lysed using a sonicator for 2 min (Branson Digital Sonifier 450, 3 mm tapered microtip, 50% amplitude, 10 s ON/ 50 s OFF) in an ice bath. Proteins were labeled using Cys-maleimide conjugation with AlexaFluor488. Recombinant BRG1-6His protein was obtained from Abcam (ab82237) and labeled with RED-tris-NTA reagent (Nanotemper MO-L018). The recombinant BRG1 protein and RED-tris-NTA were mixed 1:1 molar ratio and incubated for one hour at room temperature in PBS-T buffer (provided by Nanotemper) and used for our experiments. Fluorescently labeled RNA client ([6FAM]-UGAAGGAC) was purchased from Sigma.

**Cell culture:** HEK293T cells were grown in Dulbecco's Modified Eagle's Medium (Gibco™ 11965092) plus 10% fetal bovine serum (Gibco™ A3160501) at 37 °C and 5% $CO_2$. For transfection, 25,000 cells were seeded in 8-well chamber slides (Nunc™ Lab-Tek™ II Chambered Coverglass 155409). 36 hours later, cells were transfected with 0.5 μg plasmid using Lipofectamine 2000 reagent (Thermofisher 11668030) following the manufacturer's protocol and imaged between 36-48 hours. Lentiviral transfection were performed as described in reference [2] using Lipofectamine 2000 as the transfection reagent. A list of plasmids used for expression in HEK293T cells is listed in Table C.

**Fluorescence imaging:** For live-cell imaging, the cells were transferred to FluoroBrite DMEM Media (Thermofisher A1896701) with Hoechst33342 dye (Thermofisher H3570; 1μg/ml) an hour before imaging. The cells were incubated at 37 °C on a temperature-controlled stage and imaged with Zeiss LSM710 laser scanning confocal microscope (Plan-Apochromat 63x/1.4 oil DIC M27). For optodroplet assay, previously characterized Cry2 homo-oligomerization domain [2, 3] was fused to $FUS^{PLD}$ and $BRG1^{PLD}$ to nucleate condensation. To induce optodroplets, cells were exposed to blue light (488 nm) at 2% intensity for 1 minute while imaging. In the case of Cry2-mCherry-FUS-DDIT3 construct, the Cry2 domain was not required for the formation of FUS-DDIT3 condensates as evidenced by the condensates formed by GFP-tagged FUS-DDIT3. Post-image processing was performed with FIJI and CellProfiler [4, 5].

**Phase separation experiments:** Proteins were buffer exchanged into 25 mM Tris-HCl buffer (pH 7.5), 25 mM NaCl, and 10% Ficoll PM70 at room temperature, unless otherwise stated. The samples for experiments were prepared at the required protein concentration and buffer conditions, following which TEV protease (TEV: protein = 1:25 v/v) was added to the sample and incubated for 1 hr at 30 °C to cleave the His6-MBP-N10 tag. Samples were then placed onto a Tween20-coated (20% v/v) microscope glass slide. A custom-made containment was prepared with the broad end of a plastic pipette tip and sealed onto the glass slide. 3 μl of the sample was placed at the center and the top was sealed with parafilm to prevent evaporation. The samples were incubated for an hour and examined under a Zeiss Primo-vert inverted iLED microscope (40x or 100x objective) or the Zeiss LSM710 laser scanning confocal microscope (Plan-Apochromat 63x/1.4

oil DIC M27).For Zeiss Primovert, ZEN (blue, v2.3) was used for image recording/processing and for Zeiss LSM710, ZEN (SP5 2012 Black) was used.

**Single-molecule DNA tethering assay:** The single-molecule experiments were performed using a laser scanning confocal microscope (LUMICKS™ C-trap, 60× water-immersion objective) equipped with a microfluidics u-Flux™ system and two 1064 nm laser traps. The experiments were performed in a microfluidic glass chamber (LUMICKS™ ) with five channels that were connected to the u-Flux™ system using FEP tubings (LUMICKS™ ). Channels 1, 2, and 3 are separated by laminar flow while channels 4 and 5 are physically isolated from the rest of the flow-cell (Fig. S7). Streptavidin functionalized 4.38 μm polystyrene beads (Spherotech Inc.) were diluted to 0.01% (solids) in 10 mM PBS buffer (pH 7.5) and added to channel 1 of the microfluidic chamber. Channel 2 was filled with double-stranded lambda-phage DNA (LUMICKS™ ) biotinylated at the two ends (3' strand) at a final concentration of 20 pg/μl in 10 mM PBS buffer (pH 7.5). Channel 3 was filled with TE buffer (10 mM Tris, 1 mM EDTA, pH 7.5). Channels 1, 2, and 3 were used for DNA tethering while channels 4 and 5 were used for DNA-protein interaction assays. The proteins (FUS-DDIT3 and BRG1) were diluted to their experimental concentrations (as mentioned in respective figure legends) in TE buffer (10 mM Tris, 1 mM EDTA, pH 7.5) and flown in channels 4 and 5, respectively. All solutions (protein/DNA/buffer) were flown at high pressure until each flow-cell was filled with the desired components. Then the flow was reduced to tether a single molecule of dsDNA. Briefly, two beads were trapped in channel 1 and moved to the buffer channel (channel 3). In the buffer channel, the optical traps were calibrated under no-flow conditions (for accurate force measurements) using the power spectral analysis of the thermal fluctuations of the beads. After calibration, the beads were transferred to channel 2 (*i.e.* DNA channel) and the flow was restarted. Flow-stretched DNA sticking to one bead was tethered between two beads by moving the second bead closer to the first one in a direction opposite to the flow.The bead was moved back and forth until an increase in force was observed. The beads with the tethered DNA were then transferred to the buffer in channel 3. To confirm that we trapped a single molecule of dsDNA, we stretched the DNA by moving the first bead away from the 2nd bead while monitoring the DNA force-extension relation. If the force-extension curve for the tethered DNA overlaps with the characteristic force-extension curves of a single lambda phage dsDNA [6], then we concluded that only one DNA molecule was trapped. The single DNA tether was then transferred to the protein channel 4 containing Alexa488-labeled FUS-DDIT3. The DNA molecule was incubated and fluorescently imaged in channel 4 for 60-120 s until we observed FUS-DDIT3 condensates forming on the DNA. Next, the DNA tether with FUS-DDIT3 condensates was transferred to channel 5 containing the BRG1 protein labeled with RED-tris-NTA. The DNA with FUS-DDIT3 condensates was incubated in the BRG1 channel for 180-240 s and probed for BRG1 recruitment using confocal imaging. For confocal imaging, the 488 nm and 635 nm excitation lasers were used to image Alexa488 labeled FUS-DDIT3 and RED-tris-NTA labeled BRG1, respectively. The DNA-protein interaction experiments were performed at a constant bead-to-bead separation wherein end-to-end length of the DNA is ~16 μm.

For colocalization analysis, two intensity profiles were measured along the trapped DNA for FUS-DDIT3 (green channel) and BRG1 (red channel). At first, prominent FUS-DDIT3 condensates were selected from the peaks in FUS-DDIT3 intensity profile based on the following criteria: the intensity inside the condensate is at least 2 times the average FUS-DDIT3 intensity on the entire length of DNA. The selected peaks from the FUS-DDIT3 channel (FUS-DDIT3 condensates) were probed for overlap with BRG1 intensity peaks. In case overlap was observed and the intensity of BRG1 inside FUS-DDIT3 condensate was higher than the average BRG1 intensity across the whole DNA, the condensate was considered as BRG1-enriched. All images were analyzed using Fiji-ImageJ (version 1.52p) [2]. All intensity profiles were plotted using OriginPro (2018b). 10-12 DNA molecules were analyzed for each tested concentration of BRG1 (50 nM and 10 nM).

**Bioinformatic analysis:** *Analysis of chromatin remodeling proteins:* Amino acid sequences of all mammalian SWI/SNF complex proteins were retrieved from the UniProt database [7], by searching for all reviewed human proteins that are annotated for the gene ontology annotation "SWI/SNF superfamily-type complex" (GO:0070603). These sequences were analyzed for the presence of prion-like domains using the PLAAC prediction tool [8]. SWI/SNF components were classified into core and signature/accessory components based on the SWI/SNF Infobase [9]. Proteins that are not classified as core in the SWI/SNF Infobase were considered as accessory proteins (Fig. S4).

*Analysis of oncofusion proteins:* The list of recurrent oncofusion proteins involving transcriptional regulators was obtained from the reference [10]. Fusion junction coordinates for corresponding genes were obtained from chimerKB of ChimerDB v4.0 [11]. Fusion proteins' domain organizations were retrieved from the Pfam database [12] using AGFusion [13] (all proteins except SS18-SSX2). Domain organization for SS18-SSX2 fusion was obtained from the reference [14]. Prion domains in the oncofusion proteins were predicted by the PLAAC prediction tool [8].

**Partition coefficient analysis:** Images for partition analysis were collected using Zeiss LSM710 laser scanning confocal microscope (Plan-Apochromat 63x/1.4 oil DIC M27). Phase separated condensates were prepared as mentioned above under *"Phase separation experiments"* and incubated with fluorescently labeled proteins. All the confocal images were collected after 1 hour of sample preparation. CellProfiler was used to segment droplets and find mean intensity within each droplet ($I_{in}$), five spots for the background signal were picked manually for each image and the mean intensity of the external dilute phase was determined ($I_{out}$). The partition coefficient (k) was calculated by determining the ratio of the two intensities ($k=I_{in}/I_{out}$)

**Estimation of phase separation concentration for proteins in cells:** This analysis is based off a similar analysis performed in reference [15] . Briefly, images after one minute of light-activated optodroplet formation (or a lack thereof) were imported into CellProfiler and cells were segmented using mCherry intensity for the OptoPLD constructs (Fig. S5b). In the case of GFP-FUS-DDIT3, the protein localization was nuclear and hence the nucleus was segmented using Hoechst staining (Fig. S1F). After segmentation, mean intensity was obtained and correlated manually with the presence or absence of condensates.

**FRAP analysis:** For FRAP experiments, a circular region of interest was bleached with 100% power for ~1-2 seconds which was followed by an imaging scan for 150-300 s. The recorded Alexa488-labeled probe intensity or GFP intensity values from the bleached ROI were then corrected for photofading and normalized with respect to the pre-bleach intensities. Multiple FRAP recovery curves were averaged for each sample and the averaged curve was plotted as a function of time using OriginPro (2018b). For each time point of the FRAP recovery curve, the standard deviation of the intensity values from the three FRAP curves was taken as the uncertainty.

**Table A: List of recombinantly purified proteins used in this study**

| Protein | Purpose of the N-terminal Tag/ site-directed mutagenesis |
|---|---|
| FUS-DDIT3 | Purification |
| FUS-DDIT3 S86C | Purification/ Cys-maleimide conjugation for site-specific protein Labeling |
| FUS | Purification |
| DDIT3 | Purification |
| BRG1$^{PLD}$ | Purification/ Cys-maleimide conjugation for site-specific protein Labeling |
| BRG1$^{PLD}$ 2C | Purification/ Cys-maleimide conjugation for site-specific protein Labeling |
| ARID1A$^{PLD}$ | Purification/ Cys-maleimide conjugation for site-specific protein Labeling |
| ARID1B$^{PLD}$ | Purification/ Cys-maleimide conjugation for site-specific protein Labeling |
| SS18$^{PLD}$ 2C | Purification/ Cys-maleimide conjugation for site-specific protein Labeling |

**Table B: Primary amino acid sequences of proteins used in this study**

| Protein | Sequence |
|---|---|
| FUS-DDIT3 | MASNDYTQQATQSYGAYPTQPGQGYSQQSSQPYGQQSYSGYSQSTDTS GYGQSSYSSYGQSQNSYGTQSTPQGYGSTGGYGSSQSSQSSYGQQSSYP GYGQQPAPSSTSGSYGSSSQSSSYGQPQSGSYSQQPSYGGQQQSYGQQQ SYNPPQGYGQQNQYNSSSGGGGGGGGGVFKKEVYLHTSPHLKADVLF QTDPTAEMAAESLPFSFGTLSSWELEAWYEDLQEVLSSDENGGTYVSPP GNEEEESKIFTTLDPASLAWLTEEEPEPAEVTSTSQSPHSPDSSQSSLAQE EEEEDQGRTRKRKQSGHSPARAGKQRMKEKEQENERKVAQLAEENER LKQEIERLTREVEATRRALIDRMVNLHQA |
| FUS | MASNDYTQQATQSYGAYPTQPGQGYSQQSSQPYGQQSYSGYSQSTDTS GYGQSSYSSYGQSQNSYGTQSTPQGYGSTGGYGSSQSSQSSYGQQSSYP GYGQQPAPSSTSGSYGSSSQSSSYGQPQSGSYSQQPSYGGQQQSYGQQQ SYNPPQGYGQQNQYNSSSGGGGGGGGGNYGQDQSSMSSGGGSGGG YGNQDQSGGGGSGGYGQQDRGGRGRGGSGGGGGGGGGGYNRSSGGY EPRGRGGGRGGRGGMGGSDRGGFNKFGGPRDQGSRHDSEQDNSDNNT IFVQGLGENVTIESVADYFKQIGIIKTNKKTGQPMINLYTDRETGKLKGE ATVSFDDPPSAKAAIDWFDGKEFSGNPIKVSFATRRADFNRGGGNGRG GRGRGGPMGRGGYGGGGSGGGGRGGFPSGGGGGGGQQRAGDWKCP |

| | |
|---|---|
| | NPTCENMNFSWRNECNQCKAPKPDGPGGGPGGSHMGGNYGDDRRGG<br>RGGYDRGGYRGRGGDRGGFRGGRGGGDRGGFGPGKMDSRGEHRQDR<br>RERPY |
| DDIT3 | MELVPATPHYPADVLFQTDPTAEMACESLPFSFGTLSSWELEAWYEDL<br>QEVLSSDENGGTYVSPPGNEEEESKIFTTLDPASLAWLTEEEPEPAEVTS<br>TSQSPHSPDSSQSSLAQEEEEEDQGRTRKRKQSGHSPARAGKQRMKEKE<br>QENERKVAQLAEENERLKQEIERLTREVEATRRALIDRMVNLHQA |
| BRG1$^{PLD}$ | MSTPDPPLGGTPRPGPSPGPGPSPGAMLGPSPGPSPGSAHSMMGPSPGPP<br>SAGHPIPTQGPGGYPQDNMHQMHKPMESMHEKGMSDDPRYNQMKGM<br>GMRSGGHAGMGPPPSPMDQHSQGYPSPLGGSEHASSPVPASGPSSGPQ<br>MSSGPGGAPLDGADPQALGQQNRGPTPFNQNQLHQLRAQIMAYKMLA<br>RGQPLPDHLQMAVQGKRPMPGMQQQMPTLPPPSVSATGPGPGPGPG<br>PGPGPAPPNYSRPHGMGGPNMPPPGPSGVPPGMPGQPPGGPPKPWPEGP<br>MANAAAPTSTPQKLIPPQPTGRPSPAPPAVPPAASPVMPPQTQSPGQPAQ<br>PA |
| ARID1A$^{PLD}$ | MSNGGGGGGGAGSGGGPGAEPDLKNSNGNAGPRPALNNNLTEPPGGG<br>GGGSSDGVGAPPHSAAAALPPPAYGFGQPYGRSPSAVAAAAAAVFHQQ<br>HGGQQSPGLAALQSGGGGGLEPYAGPQQNSHDHGFPNHQYNSYYPNR<br>SAYPPPAPAYALSSPRGGTPGSGAAAAAGSKPPPSSSASASSSSSSFAQQ<br>RFGAMGGGGPSAAGGGTPQPTATPTLNQLLTSPSSARGYQGYPGGDYS<br>GGPQDGGAGKGPADMASQCWGAAAAAAAAAAAASGGAQQRSHHAPM<br>SPGSSGGGGQPLARTPQPSSPMDQMGKMRPQPYGGTNPYSQQQGPPSG<br>PQQGHGYPGQPYGSQTPQRYPMTMQGRAQSAMGGLSYTQQIPPYGQQ<br>GPSGYGQQGQTPYYNQQSPHPQQQQPPYSQQPPSQTPHAQPSYQQQPQ<br>SQPPQLQSSQPPYSQQPSQPPHQQSPAPYPSQQSTTQHPQSQPPYSQPQ<br>AQSPYQQQQPQQPAPSTLSQQAAYPQPQSQQSQQTAYSQQRFPPPQ |
| ARID1B$^{PLD}$ | MNNYYGSAAPASGGPGGRAGPCFDQHGGQQSPGMGMMHSASAAAAG<br>APGSMDPLQNSHEGYPNSQCNHYPGYSRPGAGGGGGGGGGGGGGSGG<br>GGGGGGAGAGGAGAGAVAAAAAAAAAAAAGGGGGGGYGGSSAGYGV<br>LSSPRQQGGGMMMGPGGGGAASLSKAAAGSAAGGFQRFAGQNQHPS<br>GATPTLNQLLTSPSPMMRSYGGSYPEYSSPSAPPPPPSQPQSQAAAAGA<br>AAGGQQAAAGMGLGKDMGAQYAAASPAWAAAQQRSHPAMSPGTPG<br>PTMGRSQGSPMDPMVMKRPQLYGMGSNPHSQPQQSSPYPGGSYGPPGP<br>QRYPIGIQGRTPGAMAGMQYPQQQMPPQYGQQGVSGYCQQGQQPYYS<br>QQPQPPHLPPQAQYLPSQSQQRYQPQQDMSQ |
| SS18$^{PLD}$ | MNQNMQSLLPAPPTQNMPMGPGGMNQSGPPPPPRSHNMPSDGMVGG<br>GPPAPHMQNQMNGQMPGPNHMPMQGPGPNQLNMTNSSMNMPSSSHG<br>SMGGYNHSVPSSQSMPVQNQMTMSQGQPMGNYGPRPNMSMQPNQGP<br>MMHQQPPSQQYNMPQGGGQHYQGQQPPMGMMGQVNQGNHMMGQR<br>QIPPYRPPQQGPPQQYSGQEDYYGDQYSHGGQGPPEGMNQQYYPDGH<br>NDYGYQQPSYPEQGYDRPYEDSSQHYYEGGNSQYGQQQDAYQGPPPQ<br>QGYPPQQQQYPGQQGYPGQQQGYGPSQGGPGPQYPNYPQGQGQQYG<br>GYRPTQPGPPQPPQQRPYGYDQGQYGNYQQ |

**Table C: List of plasmids used for cellular experiments**

| **Plasmid** | **Tag** | **Source plasmid** |
|---|---|---|
| GFP-FUS | mEGFP | FUS ORF (OHu24555) was cloned by Genscript into mEGFP-C1 |
| GFP-FUS-DDIT3 | mEGFP | FUS-DDIT3 was synthesized and cloned by Genscript into mEGFP-C1 |
| GFP-DDIT3 | mEGFP | DDIT3 was synthesized and cloned by Genscript into mEGFP-C1 |
| OptoBRG1$^{PLD}$ | CRY2-mCherry | BRG1$^{PLD}$ was synthesized and cloned by Genscript into pCMV-CRY2-mCherry |
| OptoFUS$^{PLD}$ [2] | CRY2-mCherry | Kind gift from Dr. Sreejith Nair |
| OptoFUS-DDIT3 | CRY2-mCherry | FUS-DDIT3 was synthesized and cloned by Genscript into pCMV-CRY2-mCherry |
| GFP-BRG1 [16] | mEGFP | Addgene:# 65391; a gift from Kyle Miller |
| GFP-BRG1$^{PLD}$ | mEGFP | BRG1$^{PLD}$ was synthesized and cloned by Genscript into mEGFP-C1 |
| mEGFP-C1 | | Addgene: #54759, a gift from Michael Davidson |
| pCMV-CRY2-mCherry [3] | | Addgene: #58368, a gift from Won Do Heo |

### Supplementary Figures

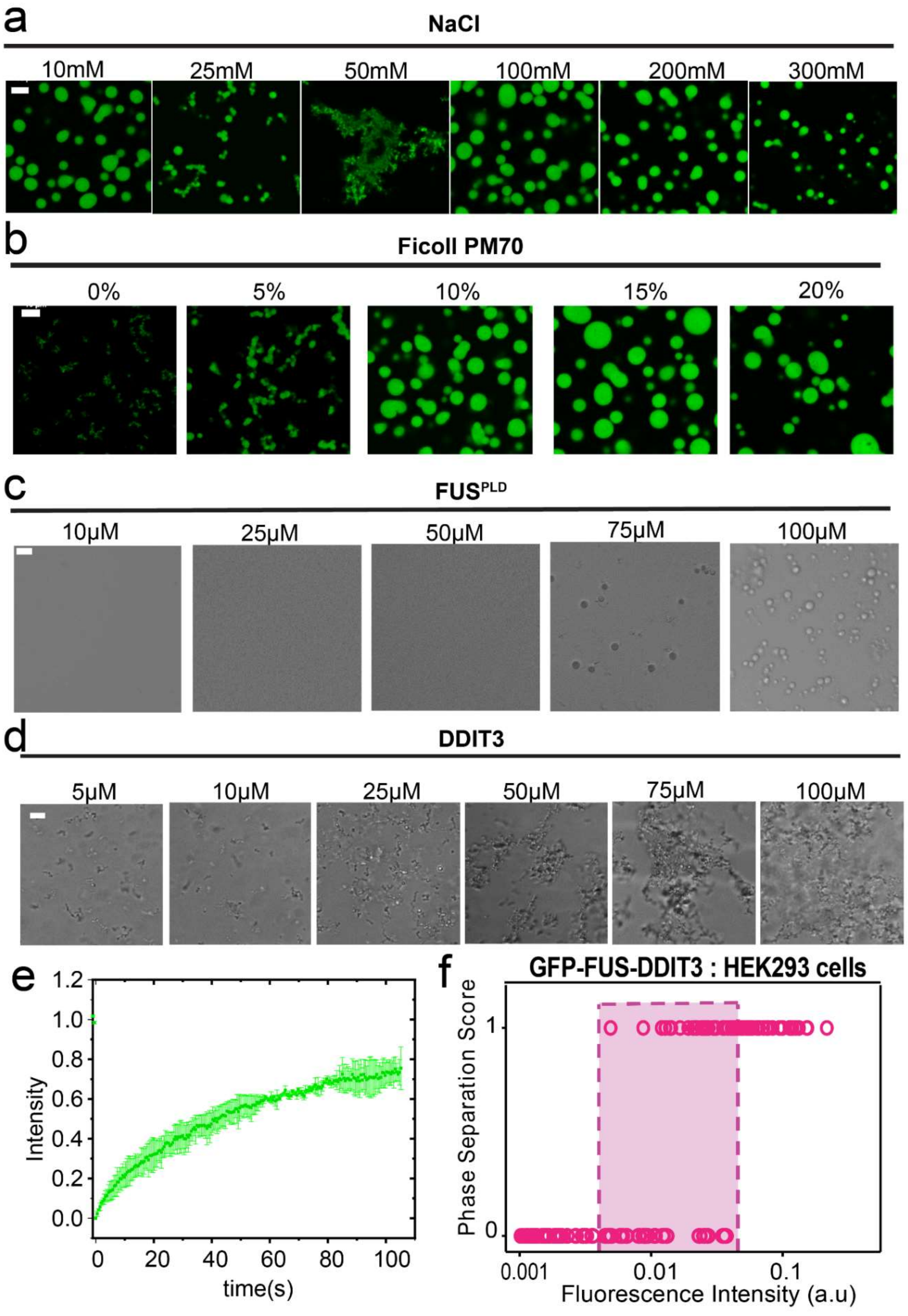

**Figure S1. Characterization of FUS-DDIT3 condensates and DDIT3 gel-like assemblies *in vitro* and in HEK293T cells.** Fluorescence microscopy images of condensates formed by the recombinantly purified FUS-DDIT3 (50 μM) containing 1% AlexaFluor488 -labeled protein with varying concentrations of **(a)** NaCl (buffer: 25 mM Tris, pH 7.5, 10% Ficoll PM70); and **(b)** Ficoll PM70 (buffer: 25 mM Tris, pH 7.5, 25 mM NaCl). **(c)** DIC images of solutions containing recombinantly purified $FUS^{PLD}$ with varying protein concentrations as indicated. **(d)** DIC images of the assemblies formed by the recombinantly purified DDIT3 with varying protein concentrations as indicated. **(e)** FRAP plots for DDIT3 condensates from *panel d* at 50 μM protein concentration. **(f)** The phase separation capacity of GFP-FUS-DDIT3 in HEK293T cells was quantified across varying levels of GFP intensity. "1" indicates the presence of condensates and "0" represents no condensates. The shaded region represents the concentration for GFP-FUS-DDIT3 in cells where a transition from no condensates to condensates was observed. The scale bar is 5 μm for all images.

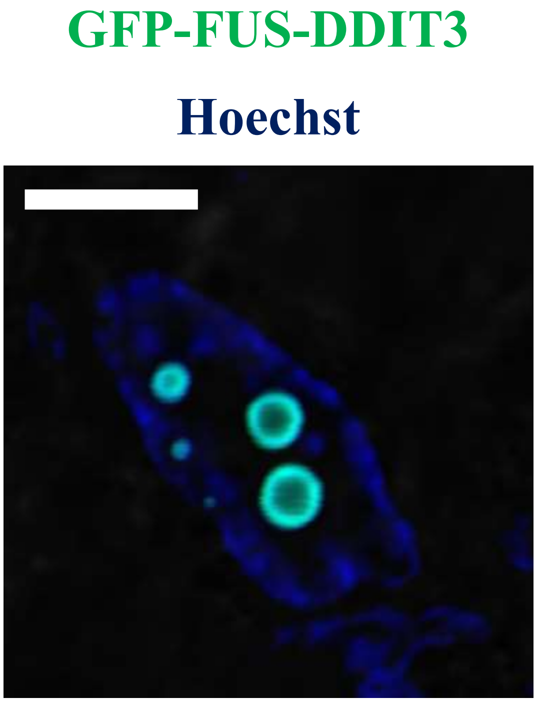


**Figure S2. A subset of GFP-FUS-DDIT3 condensates displays hollow internal space.** Fluorescence microscopy image of HEK293T cells expressing GFP-tagged FUS-DDIT3 showing a hollow morphology for FUS-DDIT3 condensates. Hoechst was used to stain the nucleus (shown in blue). The scale bar is 5 µm.

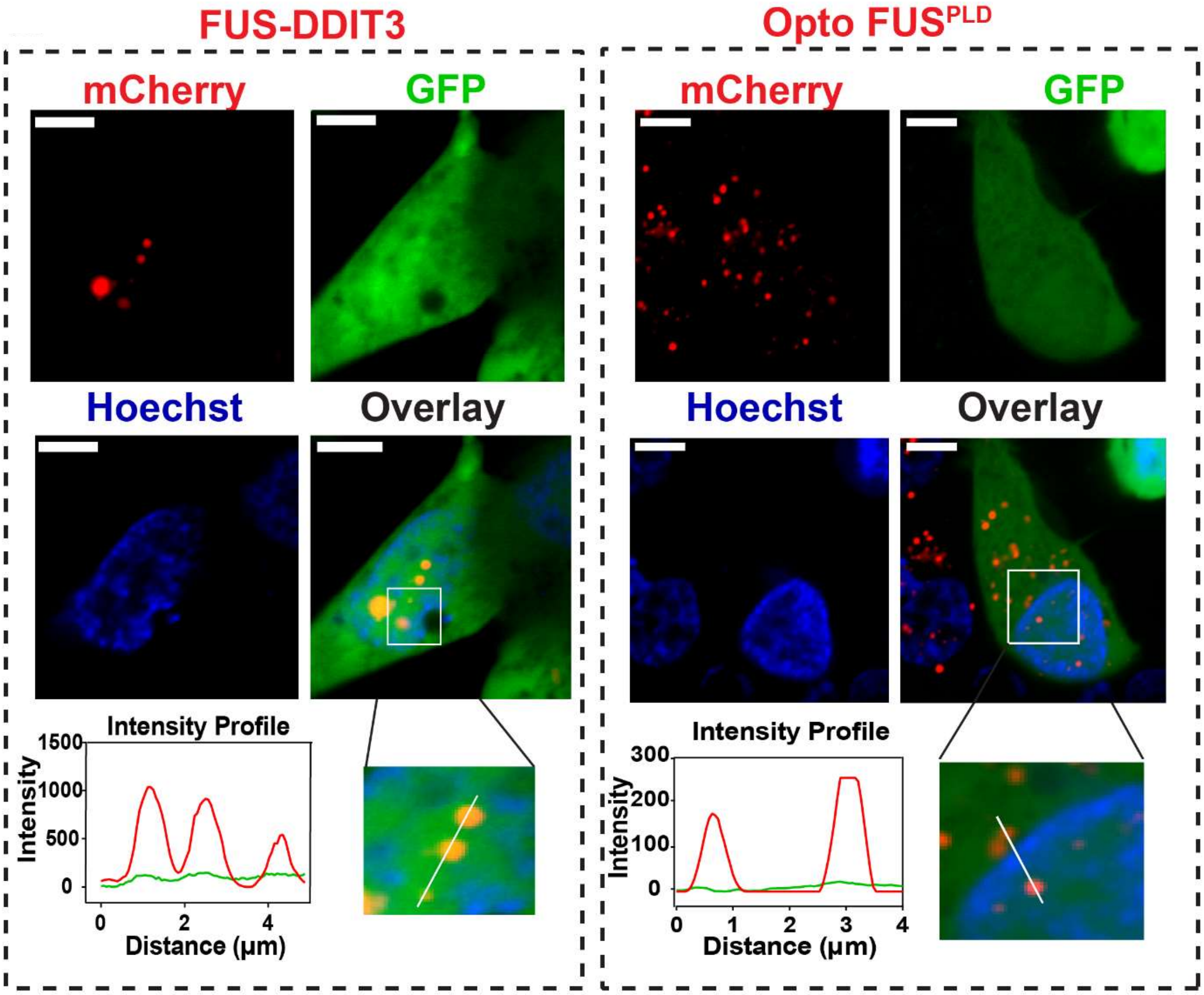


**Figure S3. GFP alone does not enrich within FUS-DDIT3 or OptoFUS$^{PLD}$ condensates.** HEK293T cells co-expressing GFP and either Cry2-mCherry-FUS-DDIT3 (*left panel*) or Cry2-mCherry-FUS$^{PLD}$: OptoFUS$^{PLD}$ (*right panel*). OptoFUS$^{PLD}$ droplets were formed by blue-light stimulation for 60 s and then enrichment of GFP was analyzed within the condensates. Cry2-mCherry-FUS-DDIT3 condensates were formed via protein overexpression without any blue-light activation. Hoechst was used to stain the nucleus. The region demarcated in the white square is magnified and the fluorescence intensity profile is shown across the linear section (white line). Green represents the intensity profile of GFP and red represents the profile for either Cry2-mCherry-FUS-DDIT3 or OptoFUS$^{PLD}$. The scale bar is 5 μm for all images.

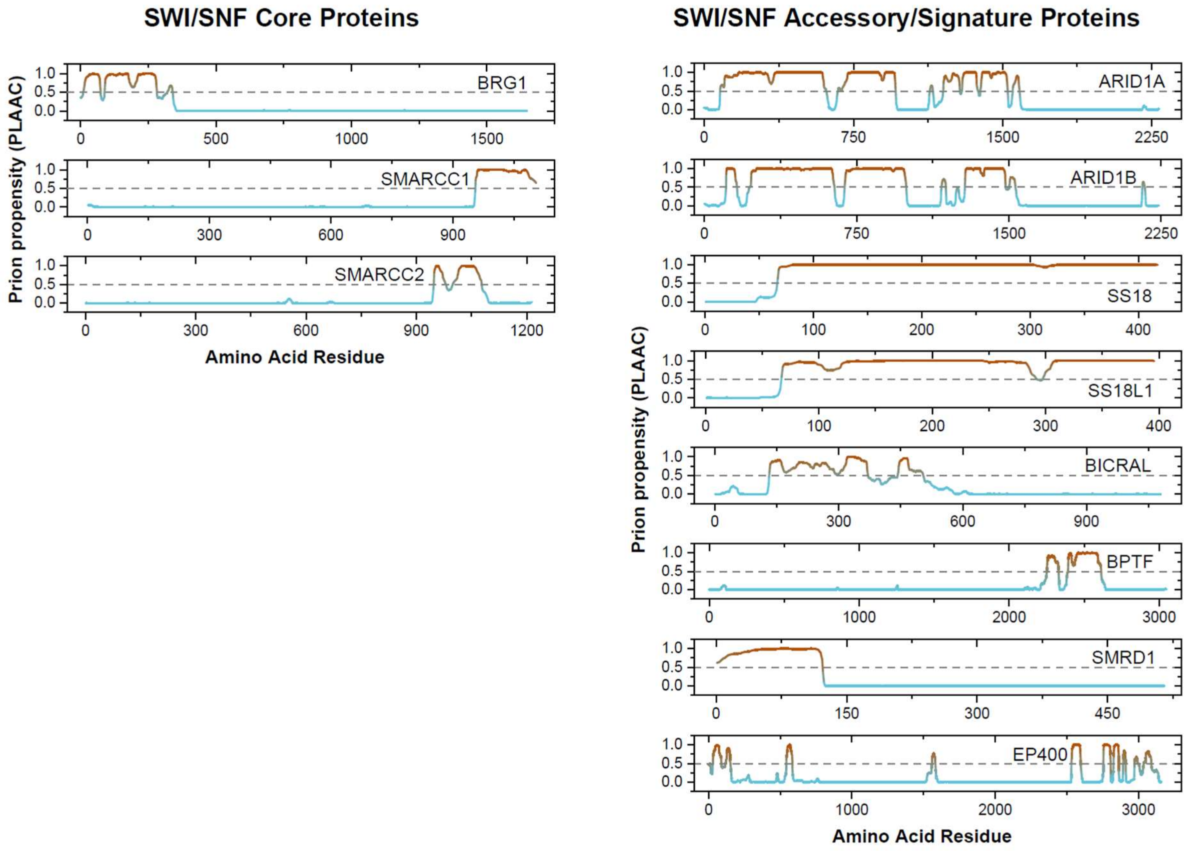


**Figure S4. Multiple subunits of the mSWI/SNF complex are enriched in prion-like domains.** PLAAC analysis showing regions of high prion-propensity for subunits of mSWI/SNF complex that showed significant prion-like domains. *Left:* PLAAC analysis for the core proteins. *Right:* PLAAC analysis of accessory proteins. Classification based on previous reports [9, 17].

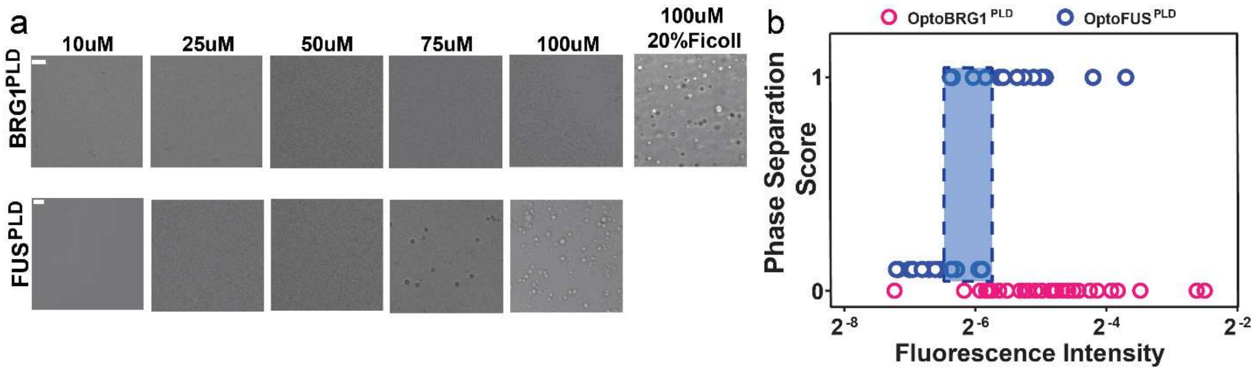


**Figure S5. The prion-like domain of BRG1 has a low intrinsic tendency to phase separate. (a)** Differential interference contrast (DIC) microscopy images of recombinantly purified BRG1$^{PLD}$ and FUS$^{PLD}$ (same as Fig. S1c) at varying protein concentrations as indicated. Condensates of BRG1$^{PLD}$ were observed only at high crowder concentrations (20% Ficoll PM70). **(b)** The phase separation capacity of Cry2-mCherry-FUS$^{PLD}$ (OptoFUS$^{PLD}$) and Cry2-mCherry-BRG1$^{PLD}$ (OptoBRG1$^{PLD}$) was quantified across varying levels of mCherry intensity in HEK293T cells. "1" indicates the presence of condensates and "0" represents no condensates. The shaded region represents the intracellular phase separation concentration for OptoFUS$^{PLD}$ from no condensates to condensates. The scale bar is 5 μm for all images.

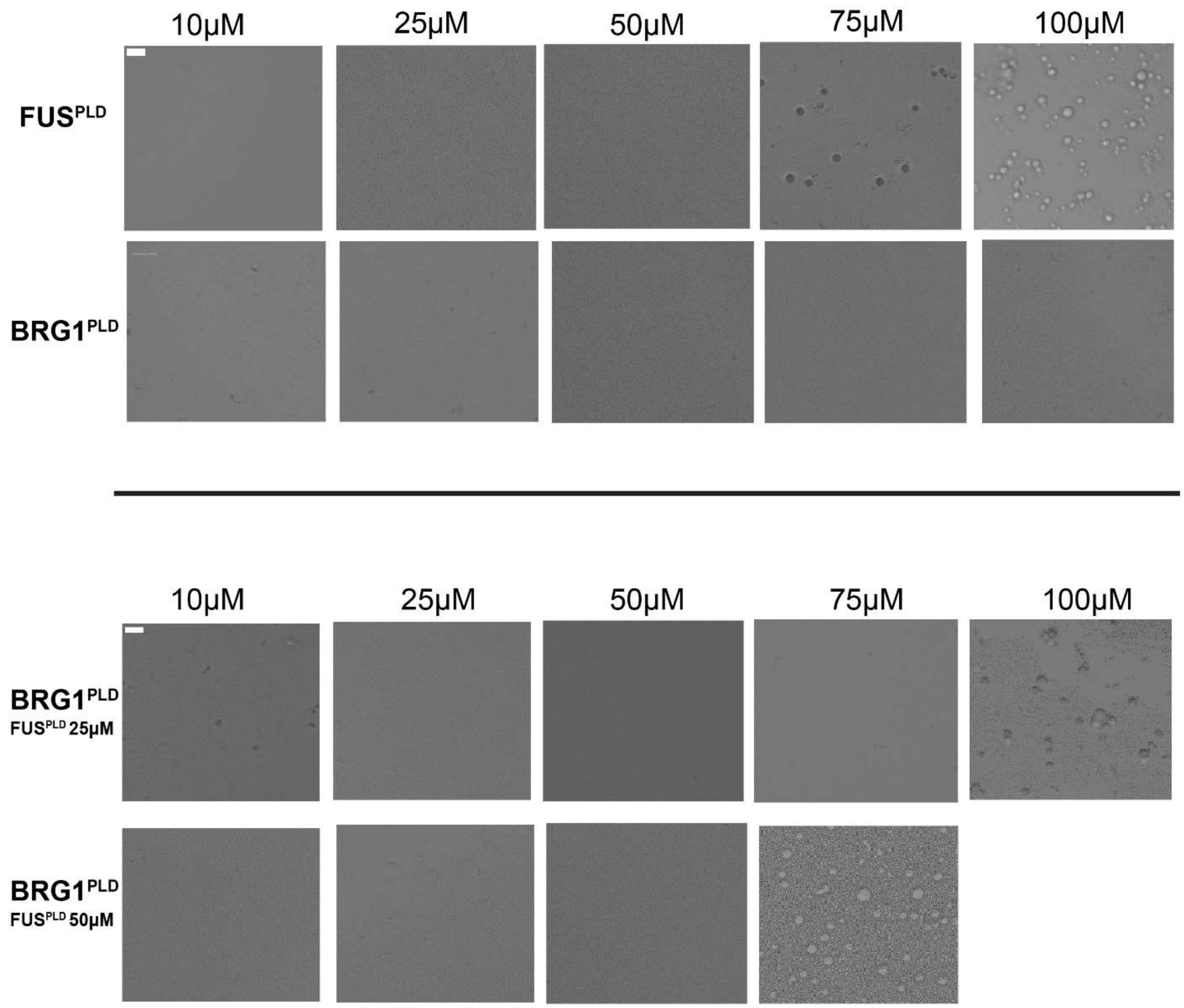


**Figure S6. Heterotypic PLD-PLD interactions drive phase separation of BRG1$^{PLD}$-FUS$^{PLD}$ mixture.** *Top panels:* Differential interference contrast (DIC) microscopy images of recombinantly purified FUS$^{PLD}$ and BRG1$^{PLD}$ at varying protein concentrations as indicated. (This panel is the same as Fig. S5a). *Bottom panels:* DIC microscopy images of recombinantly purified BRG1$^{PLD}$ mixed with either 25 µM FUS$^{PLD}$ or 50 µM FUS$^{PLD}$ at varying BRG1$^{PLD}$ concentrations as indicated. This data reveals co-condensation of the two PLDs at concentrations below their individual phase-separation threshold. The scale bar is 5 µm for all images.

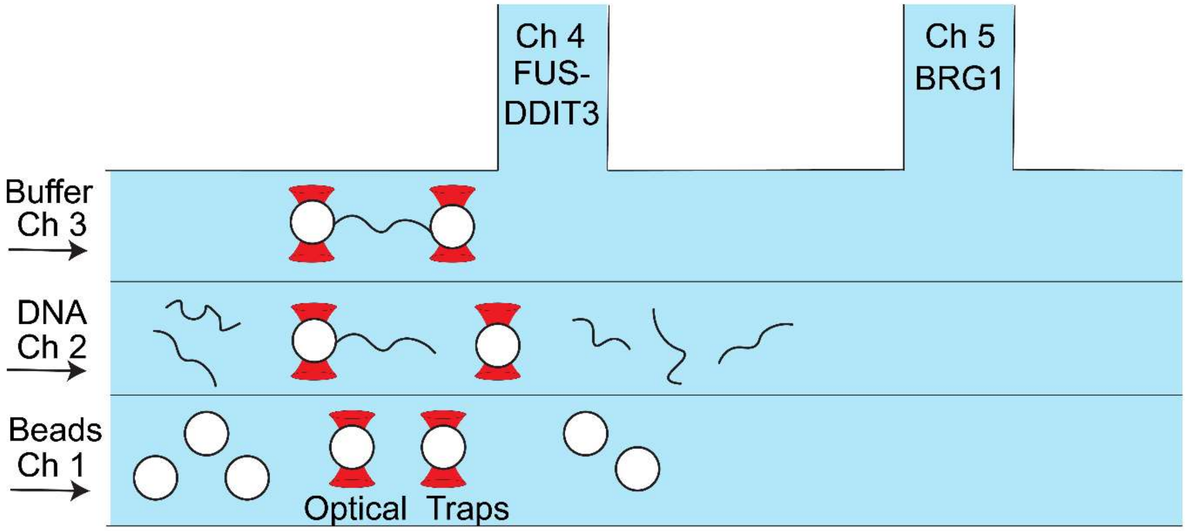


**Figure S7. Schematic of the microfluidic flow-cell used for single-molecule DNA tethering experiments.** Channels 1, 2, and 3 are separated by laminar flow while channels 4 and 5 are physically isolated from the rest of the flow-cell. Streptavidin functionalized polystyrene beads were added to channel 1 of the microfluidic chamber. Channel 2 was filled with double-stranded λ-phage DNA biotinylated at the two ends (3' strand). Channel 3 was filled with TE buffer. Channels 1, 2, and 3 were used for DNA tethering while channels 4 and 5 were used for DNA-protein interaction assays. The proteins, FUS-DDIT3 and BRG1, were flown in channels 4 and 5, respectively.

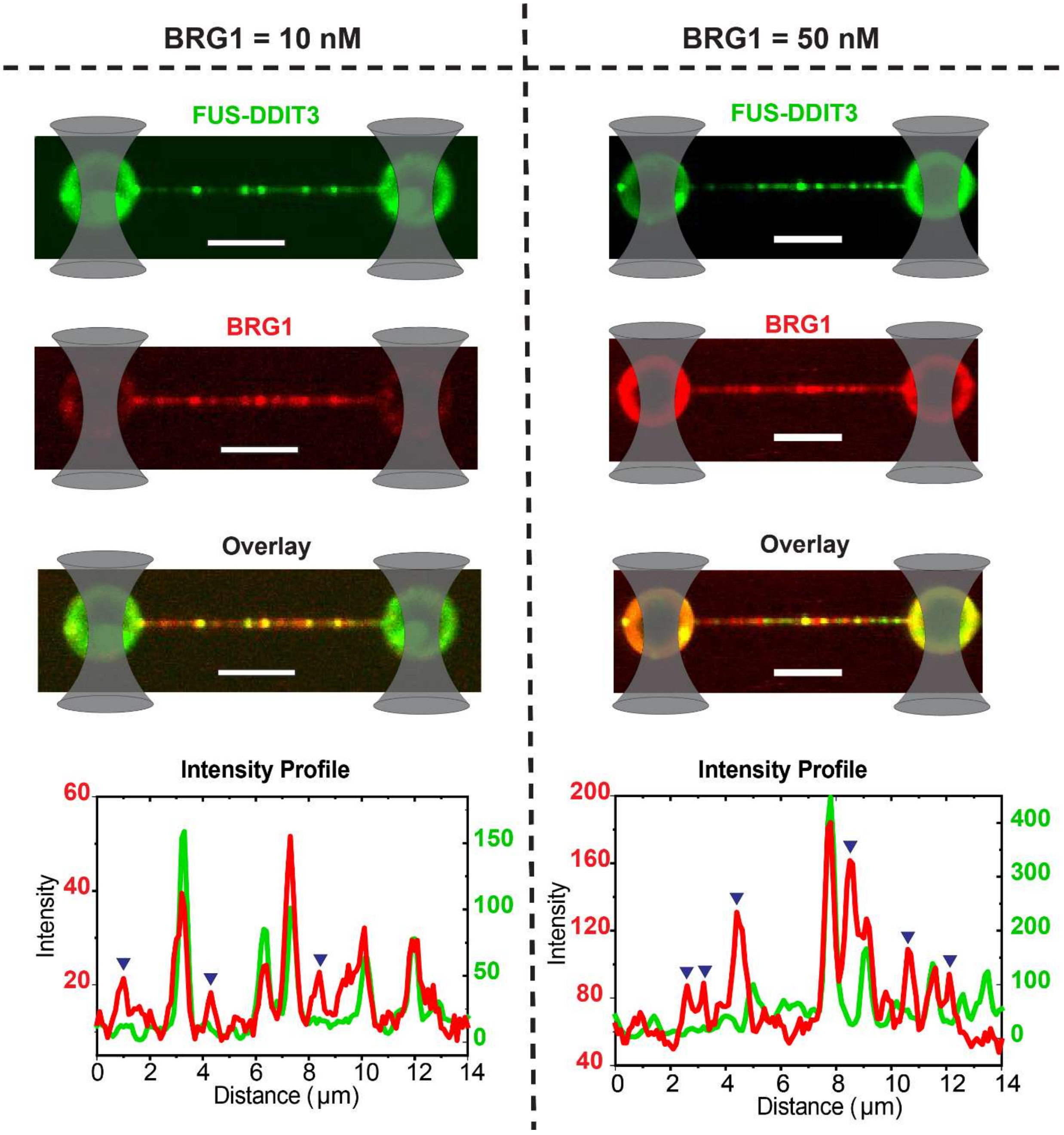


**Figure S8. BRG1 recruitment into FUS-DDIT3 condensates on DNA at different bulk BRG1 concentrations.** Multicolor confocal fluorescence micrographs and intensity profiles showing the recruitment of BRG1 (red) into the FUS-DDIT3 condensates (green) formed on a single DNA molecule at indicated concentrations of BRG1. We find that ~85% (for [BRG1] = 10 nM) and ~65% (for [BRG1] = 50 nM) of the FUS-DDIT3 foci are enriched with BRG1 (see Materials & Methods for more details). In addition, independent BRG1 peaks without a corresponding FUS-DDIT3 peak were also observed in both cases (positions marked with blue inverted triangles). [FUS-DDIT3] = 250 nM and [BRG1] = 10 nM (*left*) and [BRG1] = 50 nM (*right*). The scale bar is 5 µm for all images.
